# Synergistic insulation of regulatory domains by developmental genes and clusters of CTCF sites

**DOI:** 10.1101/2023.12.15.571760

**Authors:** Thais Ealo, Victor Sanchez-Gaya, Patricia Respuela, María Muñoz-San Martín, Elva Martin-Batista, Endika Haro, Alvaro Rada-Iglesias

**Author notes:** Equal contribution (Thais Ealo, Victor Sanchez-Gaya).

## Abstract

The specificity of gene expression during development requires the insulation of regulatory domains to avoid inappropriate enhancer-gene interactions. In vertebrates, this insulator function is mostly attributed to clusters of CTCF sites located at topologically associating domain (TAD) boundaries. However, TAD boundaries allow a certain level of physical crosstalk across regulatory domains, which is at odds with the highly specific and precise expression of developmental genes. Here we show that developmental genes and nearby clusters of CTCF sites synergistically foster the robust insulation of regulatory domains. Firstly, we found that the TADs containing developmental genes have distinctive features, including the sequential organization of developmental genes and CTCF clusters near TAD boundaries. Most importantly, by genetically dissecting representative loci in mouse embryonic stem cells, we showed that developmental genes and CTCF sites synergistically strengthened the insulation capacity of nearby boundaries through different mechanisms. Namely, while CTCF sites prevent undesirable enhancer-gene contacts (*i.e.* physical insulation), developmental genes preferentially contribute to regulatory insulation through non-structural mechanisms involving promoter competition rather than enhancer blocking. Overall, our work provides important insights into the specificity of gene regulation, which in turn might help interpreting the pathological consequences of certain structural variants.

## Introduction

The specific and precise expression of developmental genes during embryogenesis requires the concerted action of several types of *cis*- regulatory elements ^1,2^. Among them, enhancers are able to activate gene transcription by communicating with gene promoters across large linear distances ^1,3^. In contrast, insulators (also known as boundary elements) protect gene promoters from the activity of enhancers located in adjacent regulatory domains ^4,5^. In order to execute this enhancer-blocking function, insulators need to be located between an enhancer and the protected promoter. Thus, enhancer activity is typically constrained within discrete regulatory domains containing cognate gene promoters and demarcated by insulators at both ends ^4,6,7^. Insulators are best understood in *Drosophila*, where the combinatorial binding of several architectural proteins (*i.e.* Cp190, CTCF, Su(Hw), BEAF-32, GAF) to these elements enables their enhancer-blocking activity ^8–10^. In mammals, the repertoire of architectural proteins is markedly reduced and CTCF is considered the main insulator-binding protein^6,11,12^.

More recently, the emergence of novel methodologies, such as Hi-C, has resolved the 3D organization of genomes at high resolution. These studies revealed that, in both *Drosophila* and mammals, insulators often coincide with the boundaries of large contact domains that were termed “compartmental domains” or “topologically associating domains” (TADs) ^13,14^. Consequently, TADs often overlap with regulatory domains, particularly those containing major developmental genes whose expression is regulated by long-range enhancers ^15^. In mammals, the formation of TAD boundaries involves the stalling of loop-extruding cohesin complexes by CTCF ^11,16,17^. In contrast, cohesin and loop extrusion might not play a major role in TAD boundary formation in *Drosophila*, which might instead involve pairwise looping between insulator elements due to the dimerization of architectural proteins ^10,13,18^. Nevertheless, the binding of architectural proteins can only explain the formation of a fraction of TAD boundaries, suggesting that alternative mechanisms should exist ^8,10,11^. Interestingly, many architectural protein-independent boundaries are located proximal to active transcription start sites (TSS) and housekeeping genes ^14,19^. Recent reports indicate that transcription and/or active promoters can participate in the formation of contact domains and physical boundaries ^8,20–24^, but whether this can globally contribute to regulatory insulation and enhancer-blocking is largely unknown. In fact, due to the intermingling of chromatin domains, it has been shown that, at least at certain loci, the location of genes near or within boundaries might facilitate, rather than block, the communication with enhancers located along neighbouring TADs ^25–27^. Similarly, strong enhancers might be able to bypass boundaries and activate genes across TADs ^28,29^. Therefore, rather than impenetrable walls, TAD boundaries might act as dynamic and partially permeable barriers, allowing certain level of physical crosstalk across regulatory domains ^30,31^.

The permeability of TAD boundaries is at odds with the remarkable tissue specificity and spatiotemporal precision with which most developmental genes are expressed, even in the absence of CTCF ^11,18^ or other architectural proteins ^8,10,16^. Therefore, besides boundary elements, additional mechanisms might ensure the robust insulation of developmental regulatory domains. In this regard, work mostly based on the genetic dissection of the mammalian alpha and beta globin loci, as well as reporter assays in *Drosophila*, suggests that promoters can also regulate enhancer-gene communication through either promoter competition or enhancer blocking ^21,32–38^. When located within the same domain, several promoters can, in principle, share and get activated by a common enhancer ^39,40^. However, depending on the presence of distinct promoter elements, the activation of a preferred gene can preclude the expression of its neighbours independently of insulators. Promoter competition occurs when the preferred gene gets specifically activated regardless of its relative position with respect to the shared enhancer/s and neighbouring gene/s ^35^. It has been previously suggested that the competition between promoters for a shared enhancer might involve mutually exclusive enhancer-promoter contacts (“flip-flop” model) ^32,41^. However, more recent observations support non-structural mechanisms whereby promoters and enhancers share transcriptional hubs/condensates ^39,40^, within which promoters might compete for rate-limiting factors (e.g. TFs, GTFs, RNA Pol2) required for gene transcription ^42,43^. In contrast, promoter-driven enhancer blocking occurs when the preferred gene prevents the expression of its neighbours only if placed between the shared enhancer/s and the other gene/s, thus resembling how insulators work ^21,36^. Recent studies suggest that promoter-driven enhancer blocking might involve structural mechanisms, whereby protein complexes present at promoters (e.g. RNA Pol2) act as weak barriers against cohesin-mediated loop extrusion ^20,22^. It is currently unclear how prevalent these two promoter-dependent mechanisms are within endogenous loci and whether they significantly contribute to gene expression specificity, particularly in mammals. Similarly, it is largely unknown whether insulators, architectural proteins and promoters might somehow crosstalk in order to modulate the insulation of regulatory domains ^24,37^.

In the present study, we uncovered that a significant fraction of developmental genes in both mice and humans are located near TAD boundaries that contain clusters of CTCF binding sites. Furthermore, at these boundaries, developmental genes and CTCF sites are often organized in a sequential and evolutionary conserved manner, with the genes preceding the CTCF clusters. In contrast, genes previously reported as capable of bypassing boundaries ^25–27^ are typically flanked by CTCF sites and, thus, located within CTFC clusters. Most importantly, through the exhaustive genetic dissection of a couple of representative developmental loci (i.e. *Gbx2/Asb18*, *Six3/Six2*), we show that the positioning of developmental genes close to boundaries does not facilitate their own expression. Instead, we found that developmental genes and CTCF sites synergistically strengthen the regulatory insulation capacity of the nearby TAD boundaries. Finally, we show that developmental genes seem to preferentially contribute to regulatory insulation through non-structural mechanisms that involve promoter competition rather than enhancer blocking. Overall, our work provides important insights into the mechanisms contributing to the robust and specific expression of developmental genes during mammalian embryogenesis.

## Results

### TADs containing developmental genes display distinctive features

While dissecting the regulatory landscapes of a few representative developmental genes characterized by the presence of large CGI clusters and polycomb domains at their promoters (e.g. *TFAPA*, *Six3*, *Lhx5*) ^44,45^, we noticed that these genes were often located within TADs displaying rather unique features. Namely, these “developmental” TADs (i) showed low gene density and (ii) displayed a skewed gene distribution, as the developmental genes were often located near TAD boundaries (Fig. S1-2). To evaluate whether these features are prevalent among “developmental” TADs, we used TAD maps previously generated in either mouse or human cells and defined developmental genes based on the presence of broad polycomb domains around their TSS:

i. Gene density: We first classified TADs previously identified in mESC and hESC according to gene density (High, Medium or Low) and performed Gene Ontology analyses for the genes contained within each TAD category. Interestingly, TADs with low gene density were strongly and specifically enriched in developmental genes, while High and Medium gene density TADs showed milder gene ontology enrichments and were not particularly enriched in developmental terms (Fig. 1A, Fig. S3, Data S1-2). Next, to evaluate whether these observations could be generalized, we considered TAD maps previously generated in additional human (n=37) and mouse (n=14) cell types. Notably, developmental genes (i.e. genes with broad polycomb domains) were significantly enriched within TADs with low gene density in all the considered mouse and human cell types (Fisher test p-values obtained were smaller than 4.2e-05 and 1.6e-10 in humans and mice, respectively, for all tests performed, with an average odds ratio of 2.0 (95% confidence interval: 1.9 to 2.1) in humans and 2.1 in mice (95% confidence interval: 2.0 to 2.2) . In contrast, housekeeping genes were not enriched within low gene density TADs in any of the analyzed TAD maps (Fisher test p-value=1 for all tests, average odds ratio of 0.36 and 0.6 in humans and mice, respectively).
ii. Gene distribution: It was previously shown that TAD boundaries are enriched for housekeeping genes in comparison to tissue-specific genes ^46^. However, from a regulatory standpoint, developmental and tissue-specific genes represent fundamentally different gene categories, as they typically differ in the type of promoter (i.e. CpG-rich for developmental genes and CpG-poor for tissue-specific genes) and long-range enhancer responsiveness ^47,48^. To investigate the distribution of different types of genes (i.e. housekeeping, developmental, all) with respect to TADs, we divided TADs in 10 bins of equal sizes and calculated the number of genes within each bin based on the location of their TSS (Fig. 1B). In addition, genes located outside TADs (i.e. within TAD boundaries or inter-TAD genomic regions) were assigned to a bin labelled as bin 0 (Fig. 1B). Next, we computed the percentage of genes located in each bin. Notably, while housekeeping genes were often found within or close to TAD boundaries (i.e. enriched in bin 0 and bin1), developmental genes were preferentially located inside TADs and near their boundaries (i.e. enriched in bin 1 and depleted in bin 0) (Fig. 1C-E). The tendency of housekeeping genes to be located within TAD boundaries and of developmental genes to occupy intra-TAD boundary-proximal locations was more pronounced when considering high-resolution TAD maps (Fig. 1F). The preferential location of developmental genes close to TAD boundaries was similarly observed in both mice and humans, suggesting that it might represent an evolutionary conserved feature (Fig. 1C-E).

**Fig. 1:**
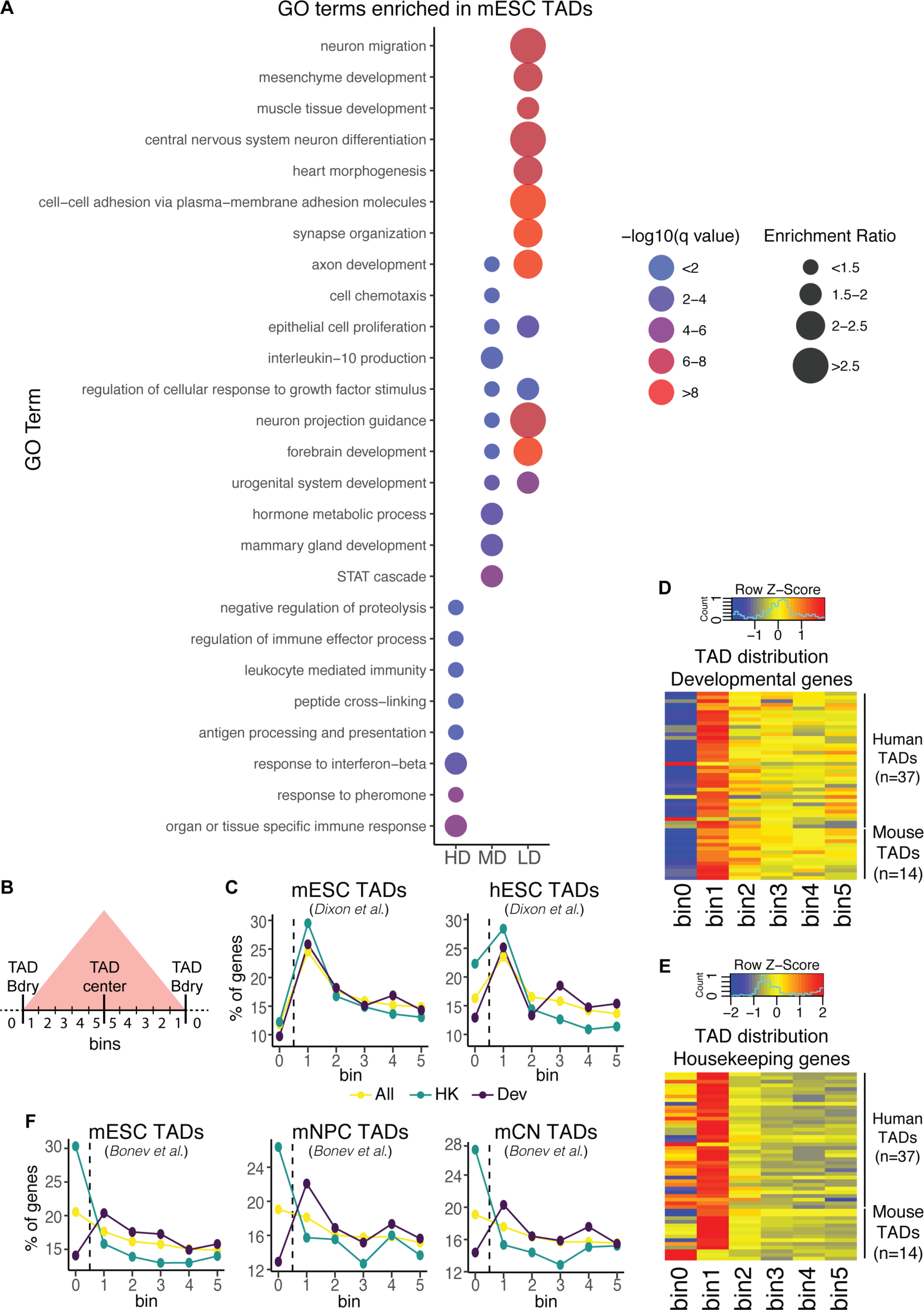
TADs containing developmental genes have distinctive features. **(A)** TADs previously identified in mESC ^14^ were classified based on their gene density in three different groups: High Density (HD), Medium Density (MD) and Low Density (LD). Then, the genes present within each TAD group were subject to GO enrichment analysis. For the MD (n=17 enriched GO terms) and LD (n=135 enriched GO terms) groups, only the top 10 most significantly enriched GO terms are shown, while for the HD group all the significantly enriched GO terms (n=8) are presented. **(B)** The distribution of different groups of genes within TADs was investigated using previously generated TAD maps. TADs were divided in 10 bins of equal sizes, which were then grouped in five bin pairs based on their distance to the nearest boundary (e.g. Bin 1 is the closest to TAD boundaries; Bin 5 is the most distal from TAD boundaries) (see Methods). Bin 0 indicates genes located at inter-TAD regions (i.e. within TAD boundaries). **(C)** Distribution of different groups of genes (All (yellow), Housekeeping (HK; green) and Developmental (Dev; purple)) within TADs according to Hi-C data previously generated in mESC ^14^ and hESC ^101^. **(D-E)** Heatmap plot (with scaling by rows) showing the distribution of developmental (D) and housekeeping (E) genes within TADs previously identified in several human (n=37) and mouse (n=14) cell types. **(F)** Distribution of different groups of genes (All (yellow), Housekeeping (HK; green) and Developmental (Dev; purple)) within TADs according to high-resolution Hi-C data previously generated in mESC, mouse neural progenitors (mNPC) and mouse cortical neurons (mCN) ^72^. In (C-F), bins were defined as described in (B).

### Sequential organization of developmental genes and clusters of CTCF sites at “developmental” boundaries

Based on the previous observations, we then evaluated whether the boundaries close to developmental genes (i.e. “developmental” boundaries) display any distinctive features. First, we compared the number of CTCF peaks and CTCF ChIP-seq aggregated signals at “developmental” and “non- developmental” (i.e. “other”) TAD boundaries (developmental boundaries are defined as those having a developmental gene located in bin 1; Fig. 1B). For both CTCF metrics, the “developmental” boundaries showed higher values (Fig. 2A-B). Interestingly, despite these differences in CTCF binding, insulation scores and boundary strength (i.e. physical insulation) at “developmental” boundaries were similar to those observed at “other” TAD boundaries (Fig. 2A-B).

**Fig. 2:**
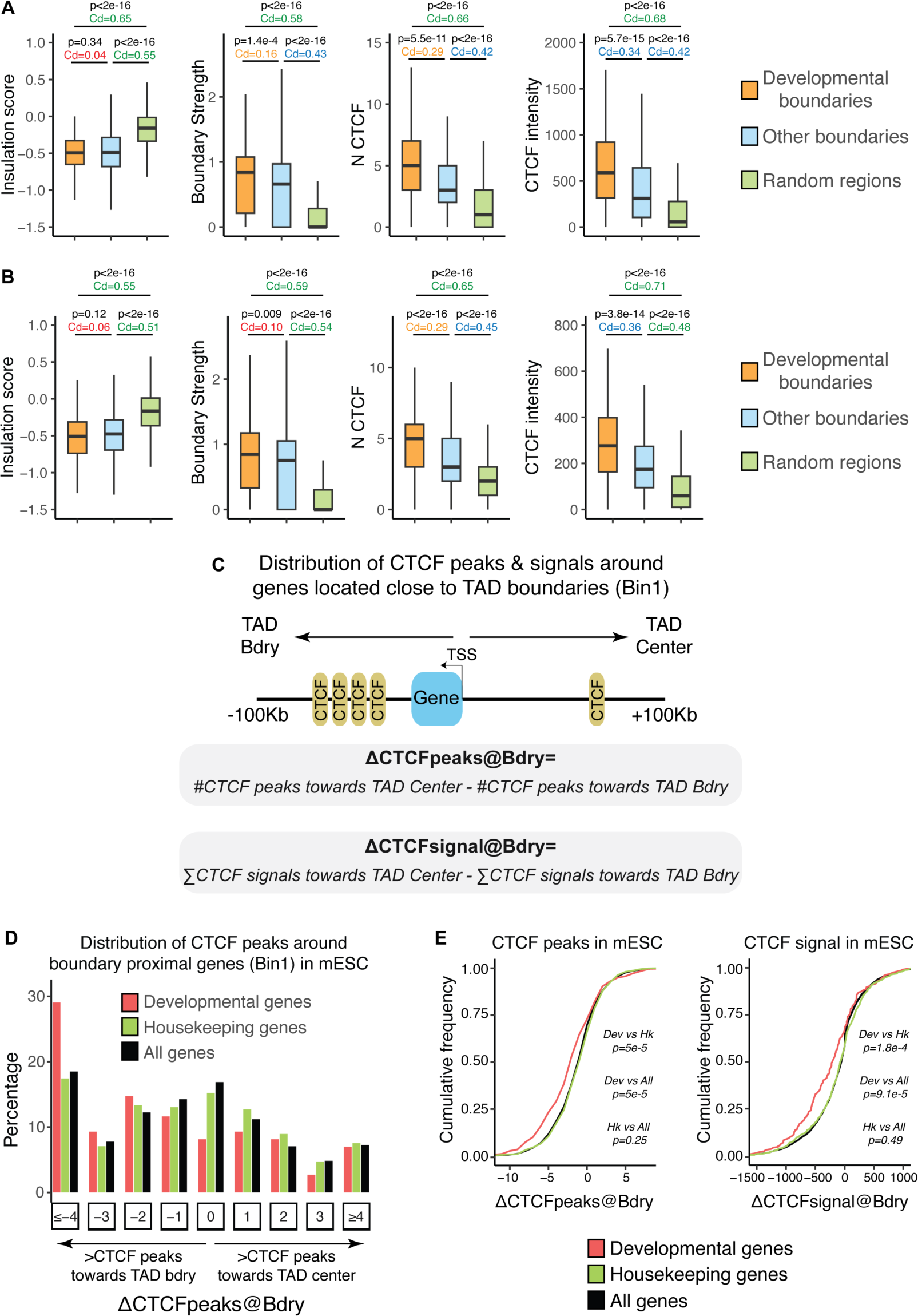
Developmental genes and clusters of CTCF sites are sequentially organized near mouse TAD boundaries. **(A-B)** TAD maps, Hi-C data and CTCF ChIP-seq profiles previously generated in either mESC (A) or hESC (B) (see Methods for details) were used to investigate (from left to right) the insulation scores, boundary strength, number of CTCF peaks and CTCF peaks aggregated signal at “developmental” TAD boundaries (n=176 for mESC, n=230 for hESC; orange), all other TAD boundaries (n=4922 for mESC, n=3618 for hESC; blue) or random regions (n=500 green). “Developmental” boundaries were defined as those having a nearby developmental gene located within bin 1 (Fig 1B). P-values were calculated using unpaired two-sided Wilcoxon tests with false discovery rate correction for multiple testing; Cliff’s delta (Cd) effect sizes are shown as coloured numbers (green: large effect size; blue: medium effect size; orange: small effect size; red: negligible effect size). **(C)** To investigate the distribution of CTCF peaks around different types of genes located close to TAD boundaries (i.e. genes located in Bin1 as described in Fig 1B), we considered the TSS of each gene as reference point and a window of +/- 100 Kb to calculate: ΔCTCFpeaks@Bdry as the difference between the number of CTCF peaks located towards the TAD center and those located towards the TAD boundary (negative values indicate that CTCF peaks are more abundant towards the TAD boundary, while positive values indicate that CTCF peaks are more abundant towards the TAD center); ΔCTCFsignal@Bdry as the difference between the aggregated signal of the CTCF peaks located towards the TAD center and the aggregated signal of the CTCF peaks located towards the TAD boundary (negative values indicate that CTCF signals are higher towards the TAD boundary, while positive values indicate that CTCF signals are higher towards the TAD center). These metrics were calculated in mESC using previously reported TAD maps and CTCF ChIP-seq profiles (see Methods). **(D)** Histogram showing the distribution of ΔCTCFpeaks@Bdry values in mESC for different types of genes (developmental, housekeeping, all) located close to TAD boundaries (bin 1 in Fig 1B). **(E)** Cumulative distribution plots for ΔCTCFpeaks@Bdry (left) and ΔCTCFsignal@Bdry (right) values in mESC show that for developmental genes the number of CTCF peaks and the CTCF signals are significantly more skewed towards negative values (i.e. the number of CTCF peaks and the associated ChIP-seq signals are higher towards TAD boundaries) than for the other considered gene categories. P-values were calculated using unpaired two-sided Wilcoxon tests with false discovery rate correction for multiple testing.

On the other hand, upon visual inspection of several “developmental” boundaries we noticed that developmental genes and CTCF clusters were sequentially organized (Fig. S1-2), with the developmental genes typically preceding most of the CTCF peaks (i.e. the CTCF peaks were preferentially located in the genomic regions separating the genes from the nearby TAD boundaries). To evaluate whether this sequential organization of developmental genes and CTCF clusters is somehow characteristic of “developmental” boundaries, we performed a global analysis of the distribution of CTCF peaks around genes located in bin 1 (Fig. 1B). Briefly, for each gene, we considered a window of +/-100 Kb around its TSS and compared the number of CTCF peaks (and associated CTCF ChIP-seq signals) located between the gene TSS and the nearby TAD boundary with the number of peaks (and associated CTCF ChIP-seq signals) located between the gene TSS and the center of its TAD (ΔCTCFpeaks@Bdry and ΔCTCFsignal@Bdry; Fig. 2C). Notably, the fraction of boundary-proximal genes in which the CTCF peaks and their associated signals were higher towards the TAD boundaries than towards the TAD centre (i.e. genes displaying negative ΔCTCFpeaks@Bdry and ΔCTCFsignal@Bdry values) was significantly larger for “developmental” genes than for other gene types (i.e. all, housekeeping) in both mice and humans (Fig. 2D-E; Fig. S4). This indicates that boundary-proximal genes in general and housekeeping genes in particular are frequently flanked by CTCF peaks, while for developmental genes the nearby CTCF peaks tend to accumulate in the genomic regions extending from the genes towards the TAD boundaries (i.e. developmental genes preceding the CTCF peaks). It has been previously proposed that the positioning of certain housekeeping (e.g. *Mcm5*) or developmental (e.g. *Dbx2, Pitx1*) genes near or within TAD boundaries might enable them to bypass those boundaries and communicate with enhancers located in two adjacent regulatory domains ^25–27^. However, upon closer inspection of some of these genes we noticed that they were flanked by CTCF sites, rather than located next to them (i.e. there is a similar number of CTCF peaks at both sides of the gene) (Fig. S5-6). Recent reports indicate that this location within, rather than adjacent to, CTCF clusters could facilitate cross-boundary contacts ^31^.

Therefore, the sequential organization of developmental genes and CTCF clusters at “developmental” boundaries seems to represent a frequent and evolutionary conserved feature of mammalian genomes. To address the potential functional relevance of this sequential organization, we genetically dissected a couple of representative loci: *Gbx2* and *Six3/Six2* (Fig. S1-2).

### The positioning of *Gbx2* near a TAD boundary does not facilitate the maintenance of its own expression in ESC

We first focused on *Gbx2*, a major developmental gene with important functions in different processes such as neural patterning or naïve pluripotency ^49,50^. *Gbx2* is highly expressed in mESC, presumably due to the regulatory activity of a super-enhancer (SE) located approximately 60 Kb downstream of its TSS (Fig. 3A). Importantly, *Gbx2* and its SE are found within a gene-poor TAD, with *Gbx2* being located ∼10 Kb upstream from three CTCF sites that constitute the 3’ TAD boundary (Fig. 3A). Moreover, this CTCF cluster separates *Gbx2* from *Asb18*, a tissue specific gene located within the neighbouring TAD (∼60 Kb downstream of the CTCF cluster) and that is inactive in mESC (Fig. 3A). To start evaluating whether the sequential positioning of *Gbx2* and the CTCF cluster at the 3’ boundary is of any functional relevance, we first generated multiple mESC clonal lines homozygous for the following re-arrangements: (i) *71Kb INV* - a 71 kb inversion that re-positions *Gbx2* and the SE with respect to the CTCF cluster, placing the enhancer in between *Gbx2* and *Asb18*; (ii) *Δ3xCTCF* - a 10 Kb deletion that eliminates the three CTCF sites, which in principle should lead to the fusion of the *Gbx2* and *Asb18* TADs; (iii) *Δ3xCTCF*:*71Kb INV* - both the 71 Kb inversion and the 10 Kb deletion (Fig. 3B; Fig. S7). Next, we measured *Gbx2* and *Asb18* expression in all the generated mESC lines (Fig. 3C-D). Regarding *Gbx2*, we found that neither the inversion nor the deletion affected its expression (Fig. 3C). This suggests that, in contrast to previous reports ^26,27,51^, neither the positioning of *Gbx2* close to the 3’ boundary nor the CTCF sites are important for the maintenance of *Gbx2* expression. In contrast, both the inversion and the deletion significantly increased *Asb18* expression, with the inversion having a larger effect (Fig. 3D). Moreover, when both re- arrangements were combined, *Asb18* expression levels were increased to a similar extent as with the inversion alone (Fig. 3D). Considering the established function of tandem CTCF sites as insulators ^52,53^, the effects of the 10 Kb deletion on *Asb18* expression were somehow expected. However, the results obtained for the inversion were more surprising and suggest that, when positioned close to the 3’ boundary, *Gbx2* might display enhancer blocking activity and, thus, contribute to the physical insulation of its own regulatory domain. To directly assess this possibility, we performed Capture-C experiments using the *Gbx2* SE or the *Asb18* promoter as viewpoints in all the previous ESC lines. As expected, the deletion of the CTCF sites reduced the physical insulation between the *Gbx2* and *Asb18* TADs, thus increasing the contact frequency between *Asb18* and the SE (Fig. 3E-F; Fig. S8). In contrast, the 71 Kb inversion strongly increased the contact frequency between the SE and the CTCF sites, but had a minor impact on the *Asb18*-SE contacts (Fig. 3E-F, Fig. S8). This suggests that the increased expression of *Asb18* in cells with the 71 Kb inversion is unlikely to be caused by the loss of *Gbx2* enhancer blocking activity. One alternative explanation is that the inversion reduces the linear distance between *Asb18* and the SE (from 141 Kb to 83 Kb), which in turn might increase *Asb18* expression without significantly affecting enhancer-gene contacts ^54–56^. In agreement with this non-linear relationship between gene activity and E-P contact frequency, the combination of the 71 Kb inversion and the CTCF deletion resulted in a strong loss of physical insulation between the *Gbx2* and *Asb18* TADs that was not translated into a further increase in *Asb18* expression compared to cells with the inversion alone (Fig. 3D-F, Fig. S8).

**Fig. 3:**
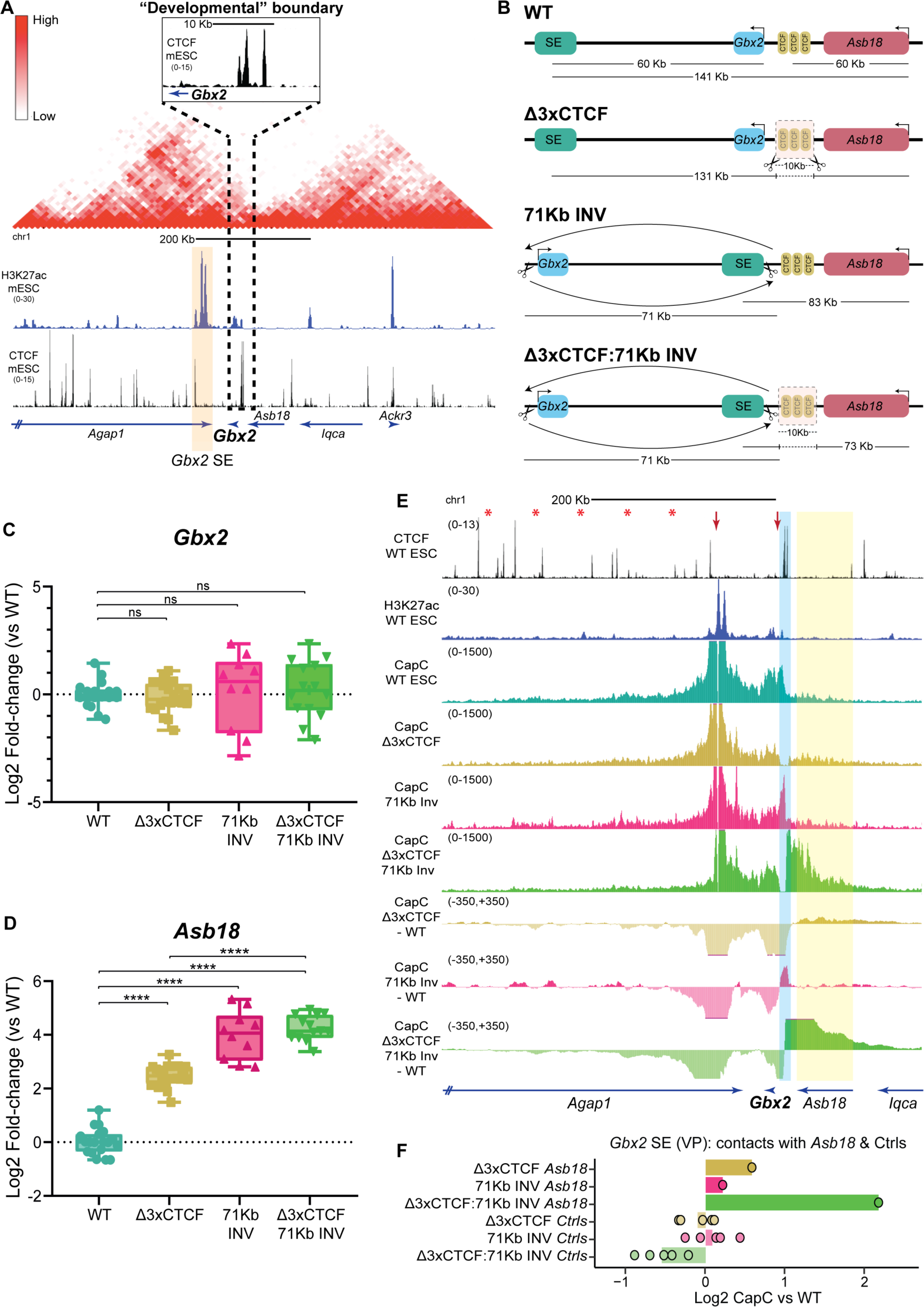
The positioning of Gbx2 close to its 3’ TAD boundary is not required to sustain its expression in mESC. **(A)** Hi-C data previously generated in mESC^72^ illustrates that Gbx2 and Asb18 are located within neighbouring TADs (i.e. Gbx2- TAD and Asb18-TAD) separated by a cluster of three CTCF sites^102^. Below the Hi-C data, H3K27ac^44^ and CTCF^102^ ChiP-seq profiles generated in mESC are also shown. **(B)** Graphical overview of different genomic rearrangements generated in mESC within the Gbx2/Asb18 locus: Δ3XCTCF - 10 Kb deletion that eliminates the three CTCF sites separating the Gbx2-TAD and Asb18-TAD; 71Kb INV - 71 kb inversion placing the Gbx2 SE close to the CTCF cluster and in between Gbx2 and Asb18; Δ3XCTCF:71Kb INV – combination of the 10 Kb deletion and 71 Kb inversion described above. The effects of these re-arrangements on the linear distances separating Gbx2, Asb18 and the SE are indicated. **(C-D)** The expression of Gbx2 (C) and Asb18 (D) was measured by RT–qPCR in ESCs that were either WT or homozygous for the genomic re-arrangements described in (B). For each cell line, Gbx2 and Asb18 expression was measured in the following number of biological replicates: WT: 18 replicates; Δ3XCTCF: 21 replicates using two different clonal lines; 71Kb INV: 10 replicates using two different clonal lines; Δ3XCTCF:71Kb INV: 14 replicates using two different clonal lines. Expression values were normalized to two housekeeping genes (Eef1a1 and Hprt1) and are presented as log2 fold- changes with respect to WT ESCs. Expression differences among ESC lines were calculated using two-sided unpaired t-tests (***fold-change > 2 and P < 0.0001; NS (not significant) fold-change < 2 or P > 0.05). **(E)** Capture-C experiments were performed as two biological replicates in WT, Δ3XCTCF, 71Kb INV and Δ3XCTCF:71Kb INV ESC using the Gbx2 SE as a viewpoint (VP). The average Capture-C signals of the two replicates performed for each mESC line are shown around the Gbx2/Asb18 locus either individually (upper tracks) or after subtracting the WT Capture-C signals (lower tracks). The TAD boundary containing the three CTCF sites is highlighted in light blue. The red arrows indicate the 71 Kb inversion breakpoints. (**F**) The average Capture-C signals shown in (E) were measured within the Asb18 gene (highlighted in yellow in (E); chr1:89952677-90014577 (mm10)) as well as within five different 30 Kb control regions (Ctrls) located within the Gbx2 TAD (red asterisks in (E) indicate the midpoint of the following controls regions (mm10): chr1:89803398-89833398; chr1:89753398-89783398; chr1:89703398-89733398; chr1:89653398-89683398; chr1:89603398-89633398). Capture-C signals are shown for the Δ3XCTCF, 71Kb INV and Δ3XCTCF:71Kb INV ESC as log2 fold-changes with respect to WT ESC.

### Robust regulatory insulation between the *Gbx2* and *Asb18* TADs depends on the synergistic effects of *Gbx2* and nearby CTCF sites

The previous results indicate that the physical insulation between the *Gbx2* and *Asb18* TADs can be largely attributed to the cluster of CTCF sites. Interestingly, despite increasing *Asb18*-SE contact frequency, the combination of the CTCF deletion and the 71 Kb inversion did not significantly change *Asb18* expression compared to the cells with the 71 Kb inversion alone (Fig. 3D-F; Fig. S8). Since promoter competition can occur regardless of the relative position of the shared enhancer(s) with respect to the competing promoters ^35,36^, this made us wonder whether, in the absence of the CTCF sites, the presence of *Gbx2* could prevent a further increase in *Asb18* expression through promoter competition rather than enhancer blocking ^35,36^. To test this hypothesis, we generated mESC lines with the following homozygous re-arrangements (Fig. S9): (i) *ΔPromGbx2* - a small (1 Kb) deletion spanning the *Gbx2* promoter that reduces *Gbx2* expression (Fig. 4A-B) and (ii) *Δ3XCTCF:ΔPromGbx2* - deletions spanning the *Gbx2* promoter and the CTCF cluster, respectively (Fig. 4A). Importantly, the 1 Kb promoter deletion should theoretically disrupt both the enhancer-blocking and promoter competition capacity of *Gbx2,* while barely changing the linear distance between *Asb18* and the SE (Fig. 4A).

**Fig. 4:**
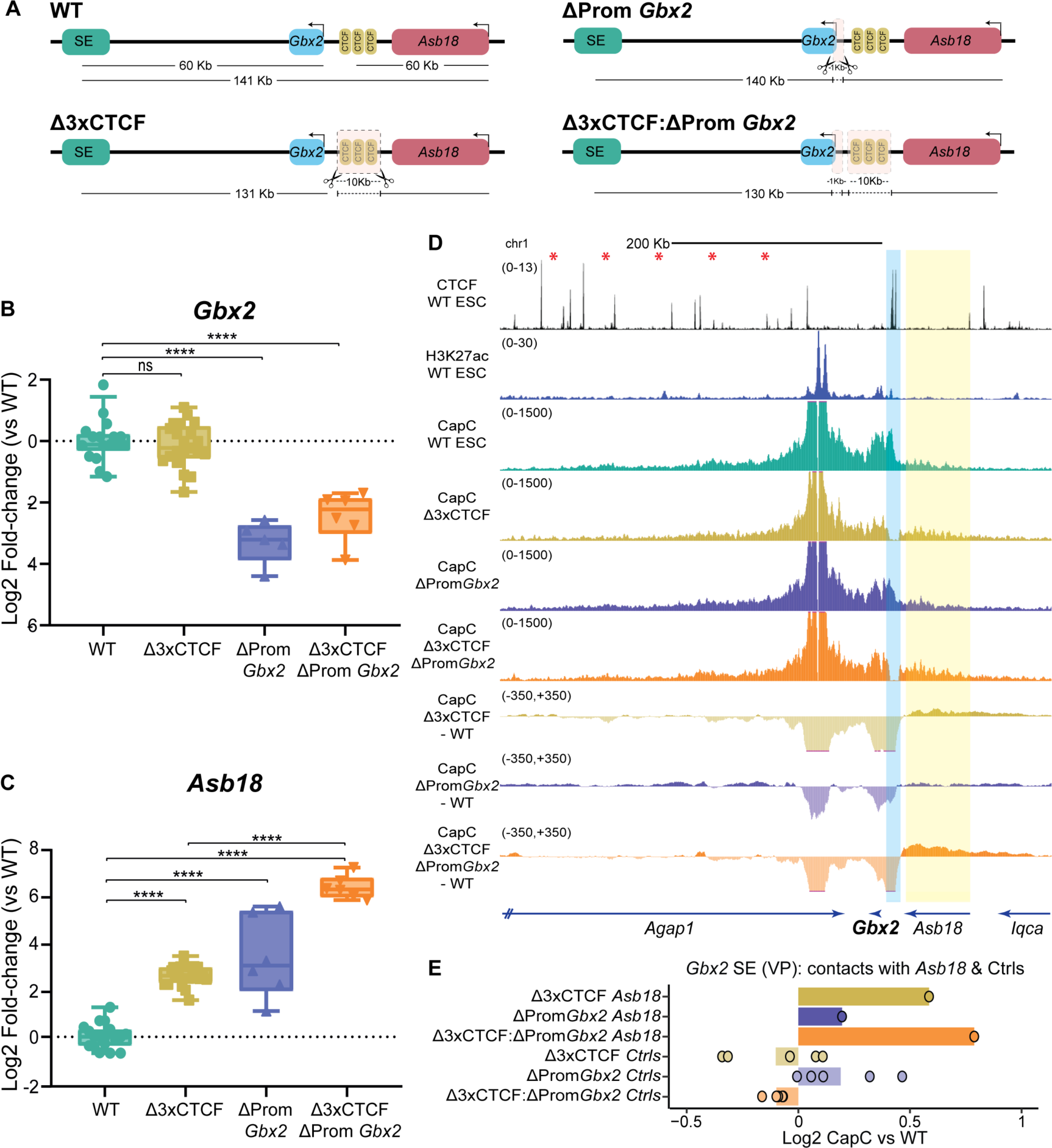
Gbx2 and the nearby CTCF cluster synergistically contribute to the regulatory insulator capacity of the Gbx2 3’ TAD boundary. **(A)** Graphical overview of different genomic rearrangements generated in mESC within the Gbx2/Asb18 locus: Δ3XCTCF - 10 Kb deletion that eliminates the three CTCF sites separating the Gbx2-TAD and Asb18-TAD (same as in Fig 3B); ΔPromGbx2 - 1 Kb deletion around the Gbx2 transcription start site (TSS) and that eliminates its main promoter; Δ3XCTCF:ΔPromGbx2 – combination of the 10 Kb and 1 Kb deletions described above. The effects of these re-arrangements on the linear distances separating Gbx2, Asb18 and the SE are indicated. **(B-C)** The expression of Gbx2 (B) and Asb18 (C) was measured by RT–qPCR in ESCs that were either WT or homozygous for the genomic re-arrangements described in (A). For each cell line, Gbx2 and Asb18 expression was measured in the following number of biological replicates: WT: 18 replicates (same data as in Fig 3C-D); Δ3XCTCF: 21 replicates using two different clonal lines (same data as in Fig 3C-D); ΔPromGbx2: 5 replicates for Gbx2 and 6 replicates for Asb18 using two different clonal lines; Δ3XCTCF:ΔPromGbx2: 6 replicates using two different clonal lines. Expression values were normalized to two housekeeping genes (Eef1a1 and Hprt1) and are presented as log2 fold-changes with respect to WT ESCs. Expression differences among ESC lines were calculated using two-sided unpaired t-tests (****fold- change > 2 and P < 0.0001; NS (not significant) fold-change < 2 or P > 0.05). **(D)** Capture-C experiments were performed as two biological replicates in WT, Δ3XCTCF, ΔPromGbx2 and Δ3XCTCF:ΔPromGbx2 ESC using the Gbx2 SE as a viewpoint. The average Capture-C signals of the two replicates performed for each mESC line are shown around the Gbx2/Asb18 locus either individually (upper tracks) or after subtracting the WT Capture-C signals (lower tracks). The three CTCF sites deleted within the Gbx2/Asb18 TAD boundary are highlighted in light blue. Capture-C tracks for WT and Δ3XCTCF ESC are the same as in Fig 3E. (E) The average Capture-C signals shown in (D) were measured within the Asb18 gene (highlighted in yellow in (D); chr1:89952677-90014577 (mm10) as well as within five different 30 Kb control regions (Ctrls) located within the Gbx2 TAD (red asterisks in (D) indicate the midpoint of the same control regions as in Fig 3F). Capture-C signals are shown for the Δ3XCTCF, ΔPromGbx2 and Δ3XCTCF:ΔPromGbx2 ESC as log2 fold-changes with respect to WT ESC.

The promoter deletion alone resulted in elevated *Asb18* expression levels (Fig. 4C). Most interestingly, the combination of the *Gbx2* promoter and CTCF cluster deletions lead to a strong and synergistic increase in *Asb18* expression (61.8 fold-change in Δ3xCTCF:ΔProm*Gbx2 vs* WT; 5.5 and 7.5 fold-changes in Δ3xCTCF and ΔProm*Gbx2 vs* WT, respectively) (Fig. 4C), resulting in *Asb18* expression levels that were considerably higher than those observed in cells with both the CTCF deletion and the 71 Kb inversion (61.8 fold-change in Δ3xCTCF;ΔProm*Gbx2 vs* WT; 17.6 fold-change in Δ3xCTCF;71 Kb INV *vs* WT) (Fig. 3D *vs* Fig. 4C). Next, we also performed Capture-C experiments in these cell lines using again the *Gbx2* SE and the *Asb18* promoters as viewpoints. Remarkably, the deletion of the *Gbx2* promoter, either alone or in combination with the CTCF cluster deletion, showed minor effects on *Asb18*-SE contact frequency in comparison to WT and Δ3xCTCF cells, respectively (Fig. 4D-E; Fig. S10). Accordingly, RAD21 ChIP-seq experiments showed that cohesin profiles within the *Asb18* locus were not significantly affected by the *Gbx2* promoter deletion (Fig. S11A). On the other hand, H3K27ac ChIP-seq signals at the *Gbx2* SE were rather similar among all the generated cell lines, albeit slightly elevated in the CTCF deletion cells (Fig. S11B), arguing that the observed *Asb18* expression changes are not caused by differences in enhancer activity.

Overall, these results support the notion of *Gbx2* preferentially contributing to regulatory (rather than physical) insulation through promoter competition. Importantly, our data also suggest that *Gbx2* and the CTCF cluster synergistically confer the nearby 3’ TAD boundary with strong insulator capacity.

### A large CTCF cluster prevents promoter competition between *Six3* and *Six2*

In order to test whether the previous observations could be extended to other “developmental” boundaries and cellular contexts, we then generated a similar set of genetic re-arrangements within the *Six3/Six2* locus (Fig. 5A). *Six3* and *Six2* are two typical developmental genes (*i.e.* with broad PcG domains and large CpG island clusters around their promoters) that are located close to each other in linear space (∼68 Kb between *Six3* and *Six2*). However, there is a strong and conserved TAD boundary containing seven CTCF sites and spanning ∼50 Kb that separates the *Six3* and *Six2* regulatory domains (Fig. 5A) ^57,58^. Accordingly, *Six3* and *Six2* display largely non-overlapping expression patterns during embryogenesis: *Six3* is expressed in the developing forebrain or the eye, while *Six2* shows high expression in the developing kidney or the facial mesenchyme ^57,58^. We previously showed that within the *Six3* TAD there is a conserved SE that specifically activates *Six3*, but not *Six2*, in neural progenitors and the developing forebrain ^44,59^. With this information at hand, we first generated mESC lines with the following homozygous re-arrangements: (i) *156Kb INV* - 156 Kb inversion between *Six3* and its SE that places the enhancer close to the TAD boundary and in between *Six3* and *Six2;* (ii) *Δ6XCTCF* - 36 Kb deletion that eliminates six out of the seven CTCF sites separating *Six3* and *Six2* ^59^; (ii) *Δ6XCTCF*:*156Kb INV* - both the 156 Kb inversion and the 36 Kb deletion (Fig. 5B; Fig. S12). Once multiple clonal mESC lines were obtained for each of these re- arrangements, we differentiated them into anterior neural progenitors (NPC) and measured the expression of *Six3* and *Six2* (Fig. 5C-D).

**Figure 5.**
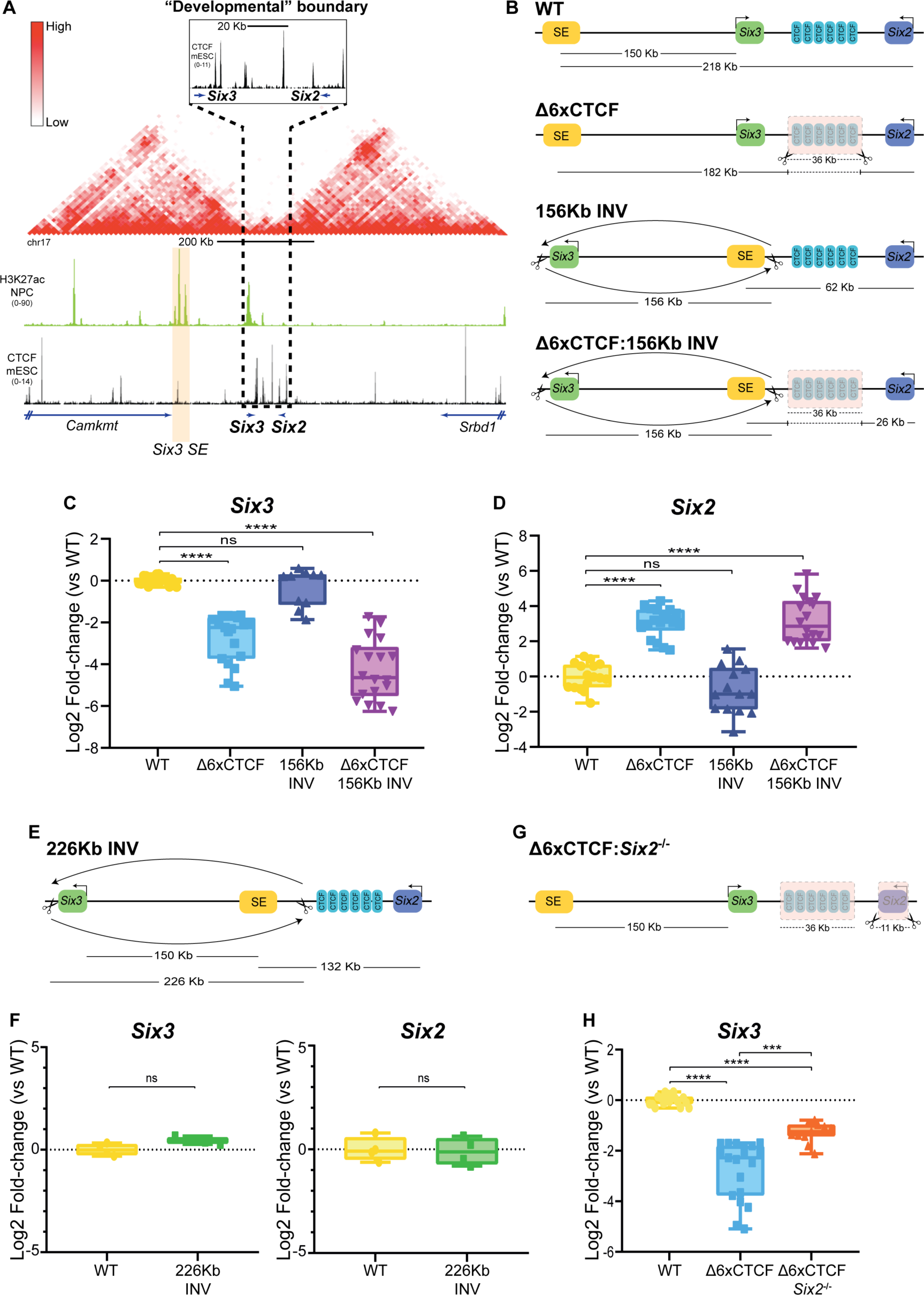
The large CTCF cluster separating the Six3 and Six2 TADs prevents promoter competition between these two genes. **(A)** Hi-C data previously generated in NPC^72^ illustrates that Six3 and Six2 are located in neighbouring TADs (i.e. Six3-TAD and Six2-TAD) separated by a cluster of seven CTCF sites^102^. Below the Hi-C data, H3K27ac^44^ and CTCF^102^ ChiP-seq profiles generated in NPC and mESC, respectively, are also shown. **(B)** Graphical overview of different genomic rearrangements generated in mESC within the Six3/Six2 locus: Δ6XCTCF - 36 Kb deletion that eliminates six out of the seven CTCF sites separating the Six3-TAD and Six2-TAD; 156Kb INV - 156 kb inversion placing the Six3 SE close to the CTCF cluster and in between Six3 and Six2; Δ6XCTCF:156Kb INV – combination of the 36 Kb deletion and 156 Kb inversion described above. The effects of these re- arrangements on the linear distances separating Six3, Six2 and the SE are indicated. **(C-D)** The expression of Six3 (C) and Six2 (D) was measured by RT–qPCR in NPC following the differentiation of ESC that were either WT or homozygous for the genomic re-arrangements described in (B). For each cell line, Six3 and Six2 expression was measured in the following number of biological replicates: WT: 16 replicates; Δ6XCTCF: 19 replicates using a previously characterized clonal line ^59^; 156Kb INV: 10 replicates for Six3 and 14 replicates for Six2 using two different clonal lines; Δ6XCTCF:156Kb INV: 18 replicates using three different clonal lines. **(E)** Graphical overview of the 226 Kb inversion generated in mESC (226Kb INV) within the Six3/Six2 locus. The effects of this inversion on the linear distances separating Six3, Six2 and the SE are indicated. (**F**) The expression of Six3 (left) and Six2 (right) was measured by RT–qPCR in NPC following the differentiation of ESC that were either WT or homozygous for the 226 Kb inversion described in (E). For each cell line, Six3 and Six2 expression was measured in the following number of biological replicates: WT: 4 replicates; 226Kb INV: 4 replicates using two different clonal lines. **(G)** Graphical overview of the 36 Kb and 11 Kb deletions generated in mESC (Δ6XCTCF:Six2^-/-^) and that eliminate the 6xCTCF cluster and Six2 gene, respectively. The effects of these deletions on the linear distances separating Six3, Six2 and the SE are indicated. (**H**) The expression of Six3 was measured by RT– qPCR in NPC following the differentiation of ESCs that were either WT or homozygous for the Δ6XCTCF and Δ6XCTCF:Six2^-/-^ deletions described above. For each cell line, Six3 was measured in the following number of biological replicates: WT: 16 replicates (same data as in C); Δ6XCTCF: 19 replicates (same data as in C); Δ6XCTCF:Six2^-/-^: 11 replicates using three different clonal lines. In C, D, F and H, the expression values were normalized to two housekeeping genes (Eef1a1 and Hprt1) and are presented as log2 fold-changes wit respect to WT NPC. The expression differences among cell lines were calculated using two-sided unpaired t-tests (****fold-change > 2 and P < 0.0001; ***fold-change > 2 and P < 0.001; NS (not significant) fold-change < 2 or P > 0.05).

Similarly to our results for the *Gbx2* locus, the 156 Kb inversion did not significantly affect *Six3* expression (Fig. 5C). It is worth noting that this inversion places *Six3* close to a single CTCF peak located next to the SE (Fig. 5A). Although we previously showed that this CTCF site was not required for *Six3* induction in NPC ^59^, it could theoretically contribute to *Six3* expression in cells with the inversion. Therefore, we engineered a 226 Kb inversion that places *Six3* further away from the TAD boundary while preserving the linear distance with respect to the enhancer (Fig. 5E; Fig. S13). Notably, this 226 Kb inversion did not affect *Six3* induction in NPC either (Fig. 5F). These results further suggest that the location of developmental genes close to TAD boundaries/CTCF clusters does not universally facilitate their own expression. Furthermore, and in contrast to our findings for the *Gbx2* locus, neither the 156 Kb nor the 226 Kb inversion affected *Six2*, which remained lowly expressed in NPC (Fig. 5D,F). This could be explained by the differences between the CTCF clusters present at each locus (seven CTCF sites spanning ∼50 Kb at the *Six3/Six2* locus *vs* three CTCF sites spanning ∼10 Kb at the *Gbx2/Asb18* locus), which is significantly larger in the *Six3/Six2* locus and, thus, could confer stronger insulation.

On the other hand, and in contrast to our observations for the *Gbx2/Asb18* locus, the deletion of the CTCF cluster not only led to the ectopic activation of *Six2* (Fig. 5D), but also significantly impaired *Six3* induction in NPC (Fig. 5C). Considering the results obtained with the inversions described above, the reduced expression of *Six3* can not be simply attributed to a potential role of the CTCF sites as facilitators of enhancer-gene communication. Alternatively, the CTCF cluster might protect *Six3* from the promoter competition activity of *Six2* and/or from the repressive effects of putative silencer elements located within the *Six2* TAD ^60^. Since there are no robust chromatin signatures to globally identify silencers, we decided to test the *Six2* promoter competition hypothesis by generating mESC lines with deletions spanning both the CTCF cluster and *Six2* (Fig. 5G; Fig. S14). Interestingly, the *Six2* deletion rescued, albeit partly, the impaired induction of *Six3* in NPC (Fig. 5H). This suggests that, rather than facilitating the communication between *Six3* and its cognate enhancers, the CTCF cluster protects *Six3* from promoter competition by *Six2* and, presumably, from silencers found within the *Six2* TAD.

### Synergistic insulation of the *Six3* and *Six2* regulatory domains by the CTCF cluster and promoter competition

The previous results suggest that, in the absence of the CTCF cluster, *Six2* competes with *Six3* for the NPC enhancers located within the *Six3* TAD. Therefore, we wondered whether this promoter competition could be reciprocal and, thus, *Six3* could participate, together with the CTCF cluster, in the robust insulation of the *Six3* regulatory domain. To explore this possibility, we initially generated a small 1 Kb deletion that removes the *Six3* promoter. However, this deletion showed a rather minor impact on *Six3* expression, probably due to the presence of alternative TSS within promoters with large CGI clusters (data not shown) ^48,61^. Therefore, we generated instead mESC lines with (i) a 27 Kb deletion spanning the *Six3* gene (i.e. *Six3^-/-^);* (ii) deletions spanning both *Six3* and the CTCF cluster (*i.e. Δ6XCTCF*:*Six3^-/-^)* (Fig. 6A; Fig. S15). All the resulting clonal ESC lines were differentiated into NPC, in which we measured *Six2* and *Six3* expression (Fig. 6A-B).

**Figure 6.**
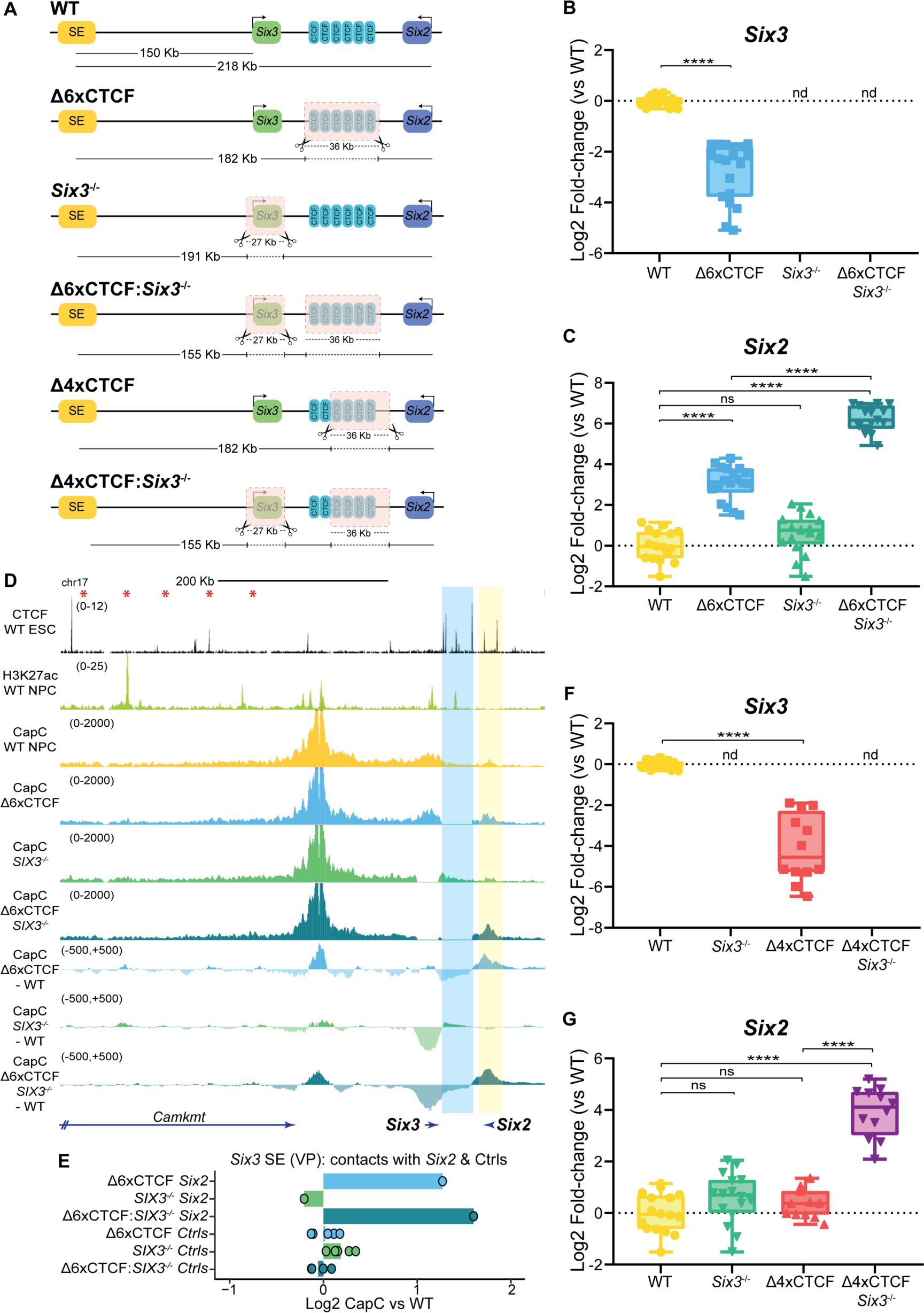
Six3 contributes to the robust insulation of the Six3 and Six2 regulatory domains through promoter competition. **(A)** Graphical overview of different genomic rearrangements generated in mESC within the Six3/Six2 locus: Δ6XCTCF - 36 Kb deletion that eliminates six out of the seven CTCF sites separating the Six3-TAD and Six2-TAD (same as in Fig 5B); Six3^-/-^ - 27 Kb deletion that eliminates the entire Six3 gene; Δ6XCTCF: Six3^-/-^ – combination of the Δ6XCTCF and Six3^-/-^ deletions described above; Δ4XCTCF – 36 Kb deletion that eliminates four out of the seven CTCF sites separating the Six3-TAD and Six2-TAD; Δ4XCTCF:Six3^-/-^ - combination of the Δ4XCTCF and Six3^-/-^ deletions described above. The effects of these re-arrangements on the linear distances separating Six3, Six2 and the SE are indicated. **(B-C)** The expression of Six3 (B) and Six2 (C) was measured by RT–qPCR in NPC following the differentiation of ESCs that were either WT or homozygous for the indicated genomic re-arrangements. For each cell line, Six3 and Six2 expression was measured in the following number of biological replicates: WT: 16 replicates (same data as in Fig 5C-D); Δ6XCTCF: 19 replicates using a previously characterized clonal line ^59^ (same data as in Fig 5C-D); Six3^-/-^ : 16 replicates using three different clonal lines; Δ6XCTCF:Six3^-/-^ : 15 replicates using three different clonal lines. **(D)** Capture-C experiments were performed as two biological replicates in WT, Δ6XCTCF, Six3^-/-^ and Δ6XCTCF:Six3^-/-^ NPC using the Six3 SE as a viewpoint. The average Capture-C signals of the two replicates performed for each cell line are shown around the Six3/Six2 locus either individually (upper tracks) or after subtracting the WT Capture-C signals (lower tracks). The six CTCF sites deleted within the Six3/Six2 TAD boundary are highlighted in blue. (**E**) The average Capture-C signals shown in (D) were measured around Six2 (highlighted in yellow in (D); chr17:85674267-85698254 (mm10)) as well as within five different 30 Kb control regions (Ctrls) located within the Six3 TAD (red asterisks in (D) indicate the midpoint of the following controls regions (mm10): chr17:85388601-85418601; chr17:85338601-85368601, chr17:85288601-85318601, chr17:85238601-85268601, chr17:85188601-85218601). Capture-C signals are shown for the Δ6XCTCF, Six3^-/-^ and Δ6XCTCF:Six3^-/-^ NPC as log2 fold-changes with respect to WT NPC. **(F-G)** The expression of Six3 (F) and Six2 (G) was measured by RT–qPCR in NPC following the differentiation of ESCs that were either WT or homozygous for the indicated genomic re-arrangements. For each cell line, Six3 and Six2 expression was measured in the following number of biological replicates: WT: 16 replicates (same data as in Fig 5C-D); Six3^-/-^: 16 replicates using three different clonal lines (same data as in Fig 5C-D); Δ4XCTCF: 12 replicates for Six3 and 11 replicates for Six2 using three different clonal lines; Δ4XCTCF:Six3^-/-^: 12 replicates using three different clonal lines. In B, C, F and G, the expression values were normalized to two housekeeping genes (Eef1a1 and Hprt1) and are presented as log2 fold-changes with respect to WT NPC. The expression differences among cell lines were calculated using two-sided non-paired t-tests (****fold-change > 2 and P < 0.0001; NS (not significant) fold-change < 2 or P > 0.05); ND (not detectable).

Remarkably, although the *Six3* deletion alone mildly increased (albeit in a non-statistically significant manner) *Six2* expression in NPC, *Six2* expression was strongly and synergistically augmented when the *Six3* and CTCF cluster deletions were combined (Fig. 6C) (72.1 Fold-change in Δ6xCTCF:*Six3^-/-^ vs* WT ; 8.8 fold change in Δ6xCTCF and 1.7 fold-change in *Six3^-/-^ vs* WT, respectively), thus in agreement with the contribution of *Six3* to regulatory insulation. Next, we performed Capture-C experiments in these cell lines using the *Six3* SE or the *Six2* promoter as viewpoints. As expected, the deletion of the CTCF cluster led to a loss of insulation between the *Six3* and *Six2* TADs and increased the contact frequency between *Six2* and the SE (Fig. 6D-E; Fig. S16). In addition, this deletion strongly increased the contacts between *Six3* and *Six2,* in agreement with the formation of a shared transcriptional hub (Fig. S16A) ^39^. Notably, the deletion of *Six3* did not change the *Six2*-SE contact frequency in comparison to WT cells (Fig. 6D-E; Fig. S16). Furthermore, the combined deletion of *Six3* and the CTCF cluster, which resulted in a strong induction of *Six2*, moderately increased the *Six2*- SE contact frequency in comparison to the CTCF deletion alone (Fig. 6D-E; Fig. S16). These results suggest that, as observed for the *Gbx2/Asb18* locus, *Six3* preferentially contributes to regulatory insulation through promoter competition, while structural mechanisms, such as enhancer blocking, seem to have a comparably smaller, albeit non-negligible, contribution. Importantly, the contribution of *Six3* to regulatory insulation through promoter competition can also explain why the 156Kb and 226Kb inversions described above did not have any major impact on *Six2* expression either alone or in combination with the CTCF cluster deletion (Fig. 5D,F). In agreement with this model, RAD21 ChIP-seq experiments showed that the CTCF cluster deletion increased cohesin levels around *Six2*, particularly at a CTCF site located upstream of this gene (red asterisk in Fig. S17A), while cohesin profiles were not affected by the *Six3* deletion (Fig. S17A). On the other hand, H3K27ac ChIP-seq signals at the *Six3* SE were rather similar among all the generated cell lines, except in the *Six3^-/-^*cells in which we observed higher H3K27ac levels, arguing that the observed *Six2* expression changes are not caused by differences in enhancer activity (Fig. S17B).

Overall, our data suggest that, as for the *Gbx2/Asb18* locus, *Six3* and the CTCF cluster synergistically contribute to the robust insulation of the *Six3* and *Six2* regulatory domains.

### Large CTCF clusters prevent competition between adjacent developmental genes

One major difference between the results obtained for the *Gbx2/Asb18* and *Six3/Six2* loci is that the CTCF cluster separating the *Six3* and *Six2* domains protects *Six3* from the repressive effects of *Six2* and, potentially, other regulatory elements within its TAD. In contrast, *Gbx2* expression was not affected by the deletion of the nearby CTCF cluster, suggesting that *Asb18* does not significantly compete with *Gbx2* for the ESC SE. This could be attributed to the fact that *Six2* and *Six3* are both typical developmental genes, which are characterized by having promoters with large clusters of CpG islands and strong long-range enhancer responsiveness (Fig. 5A) ^47,48,59^. On the contrary, while *Gbx2* is also a developmental gene with a large CGI cluster around its promoter, *Asb18* is a tissue specific gene with a CpG-poor promoter, which typically shows weak long-range enhancer responsiveness (Fig. 3A) ^47,59^. Consequently, in the absence of CTCF sites, *Six2* and *Six3* might engage into strong promoter competition for shared enhancers, while *Asb18* might represent a weak competitor against *Gbx2*. Furthermore, the regulatory domains of developmental genes, such as *Six2*, might often contain silencers whose long-range repressive capacity needs to be properly insulated ^60,62^. Therefore, we hypothesize that the boundary insulating the *Six3* and *Six2* regulatory domains might require a particularly large CTCF cluster (seven CTCF sites spanning ∼50 Kb) in comparison to the one found between *Gbx2* and *Asb18* (three CTCF sites spanning ∼10 Kb) (Fig. 3A, Fig. 5A). To test this prediction, we generated mESC lines with a deletion that removed four out of the seven CTCF sites separating the *Six3* and *Six2* regulatory domains (Fig. 6A; Fig. S18). In addition, we also generated mESC lines with the partial deletion of the CTCF cluster plus the *Six3* deletion (Fig. 6A). Interestingly, the partial deletion of the CTCF cluster alone did not significantly increase *Six2* expression, but strongly impaired *Six3* induction (Fig. 6F-G). This further suggests that, rather than facilitating gene expression, large CTCF clusters can often protect developmental genes from the repressive effects of other neighbouring developmental genes and their associated silencers ^28,60^. Last but not least, the combination of the partial deletion of the CTCF cluster and the *Six3* deletion led to a strong and synergistic induction of *Six2* in NPC (Fig. 6G). This further supports that the robust insulation provided by “developmental” boundaries requires the synergistic contribution of CTCF clusters and transcriptionally active developmental genes.

## Discussion

During embryogenesis, developmental genes are typically expressed in several spatial and/or temporal contexts, which is enabled by the presence of multiple, distinct and highly specific enhancers within regulatory domains delimited by insulators (e.g. CTCF sites) ^3,63^. Moreover, within a particular cellular context, the expression of developmental genes is often controlled by enhancer clusters (i.e. SE, locus control regions (LCR)) with strong regulatory activity ^64^. However, CTCF sites might not be sufficient to restrain the regulatory activity of these complex enhancer landscapes towards their target genes ^28,29^, suggesting that additional mechanisms might be necessary to ensure gene expression specificity and the robust insulation of “developmental” TADs. Here we report that “developmental” TADs are gene-poor and often delimited by rather unique boundaries. More specifically, we show that, within these TADs, developmental genes and CTCF clusters are often sequentially organized (with CTCF peaks being predominantly located between genes and TAD boundaries) and synergistically strengthen the regulatory insulation capacity of nearby boundaries, thus contributing to the establishment of specific expression patterns during development.

The large size and low gene density of “developmental” TADs might be required to accommodate the complex enhancer landscapes responsible for the specific, yet diverse, spatiotemporal expression patterns that many developmental genes display during embryogenesis ^3,63,65^. Furthermore, due to the long-range responsiveness of developmental gene promoters ^47,59^, their cognate enhancers can be separated by large linear distances and, thus, located anywhere within “developmental” TADs. In contrast, the regulatory elements controlling the expression of either housekeeping or tissue-specific genes tend to be proximal or even embedded within promoter regions ^66,67^. Moreover, as housekeeping and tissue-specific genes show lower long-range enhancer responsiveness than developmental genes, they might be able to co-exist with other genes with similar promoter types within gene-rich TADs without interfering with each other’s expression profiles ^15,68^.

Here we show that developmental genes and CTCF clusters synergistically strengthen the insulation capacity of nearby boundaries (i.e. “developmental” boundaries). In agreement with previous work, the physical insulation of the investigated TADs was mostly dependent on the CTCF clusters ^31,52,53,69^. In contrast, the investigated developmental genes do not seem to act as strong physical barriers preventing ectopic enhancer-gene communication (*i.e.* enhancer blocking), but preferentially contribute to regulatory insulation through promoter competition. The contribution of developmental genes to regulatory insulation through non-structural mechanisms is also in agreement with the fact that “developmental” boundaries show similar insulation scores (*i.e.* physical insulation) to those observed for TAD boundaries in general (Fig. 2A-B). We hypothesize that, by being close to boundaries, developmental genes are more likely to be located at shorter linear distances from their cognate enhancers than the potential competing genes located at the other side of the boundaries. Considering that short linear distances facilitate the functional communication between genes and enhancers ^55,56,70^, this might give developmental genes a competitive advantage over genes located on the other side of the “developmental” boundary ^71^. At the molecular level, it is currently unclear which mechanisms might explain how active genes contribute to insulation through promoter competition. Recent work suggests that RNA Pol2 can act as a weak physical barrier against cohesin-mediated loop extrusion ^20,22^. However, forced activation of candidate genes in ESC does not cause changes in chromatin insulation, suggesting that transcription and RNA Pol2 recruitment are not sufficient for the emergence of TAD boundaries ^72^. Moreover, the potential role of RNA Pol2 as a loop extrusion barrier can not explain how developmental genes contribute to insulation through promoter competition, as this does not require the placement of developmental genes in between enhancers and non-target genes (Fig. 3 and 5) ^35,36^. Instead, promoter competition might involve non-structural mechanisms whereby multiple promoters compete for rate-limiting factors (e.g. TFs, GTFs, RNA Pol2) present within shared transcriptional hubs ^42,43,73^. Although these models and the putative rate-limiting factors await experimental validation, we speculate that nearby promoters with strong enhancer responsiveness and similar architecture (e.g. *Six3* and *Six2*) might be particularly sensitive to promoter competition unless insulated by large CTCF clusters. On the other hand, it is also possible that the investigated developmental genes can act as weak enhancer blockers whose effects on physical insulation can not be easily detected due to the current resolution and sensitivity of Capture-C and other 3C methods, but that, nevertheless, can still lead to strong transcriptional changes ^55,74^. Furthermore, in the context of boundaries with no or weak CTCF sites, the contribution of developmental genes to physical insulation might be larger ^21,32–34^. In those cases, the positioning of developmental genes close to the boundaries should often place them in between their cognate enhancers and non-target genes located in the adjacent domain, thus enabling developmental genes to display enhancer blocking function.

It has been previously proposed that TAD boundaries and the associated CTCF sites can (i) facilitate the communication between genes and enhancers located within the same domain and (ii) insulate genes from the regulatory activity of enhancers located in other domains. Although CTCF sites might facilitate ultra long-range enhancer-gene contacts (e.g. *Shh, Sox9*) ^75,76^, at shorter distances CTCF sites seem to be dispensable and other types of tethering elements (e.g. CpG islands in vertebrates, GAGA elements in *Drosophila*) can mediate intra-TAD enhancer-gene communication ^59,60^. Furthermore, our data as well as recent work in *Drosophila* ^60^ suggest that, in the context of “developmental” boundaries, the main role of architectural protein clusters (e.g. CTCF) is to robustly insulate the regulatory domains of major developmental genes. Accordingly, global 3D genome organization studies show that developmental genes and their cognate enhancers are typically located within the same TADs, suggesting a strong functional overlap between topological and regulatory domains ^15^. Nevertheless, there are loci, with the *Hox* gene clusters being a notable example, in which genes and their cognate enhancers can be located in different TADs ^25,27,77^. In these cases, the location of genes within or close to TAD boundaries might allow them to bypass those boundaries and communicate with enhancers located in more than one TAD in order to achieve proper gene expression patterns ^26^. Upon closer inspection of genes capable of bypassing boundaries (Fig. S5-6), we noticed that they are typically flanked by strong CTCF sites and, thus, located within, rather than next to, CTCF clusters. This organization might favour boundary bypass, TAD intermingling and communication with regulatory elements located on either side of the genes ^26,27,31,78^. In contrast, for the two developmental genes that we investigated, we found no evidences indicating that CTCF clusters facilitate their expression or the communication with their cognate enhancers. Instead, we found that the sequential organization of genes and CTCF clusters at “developmental” boundaries is a prevalent and evolutionary conserved feature that seems to strengthen the insulation of important developmental loci.

Lastly, our findings might also have relevant medical implications, particularly in the context of structural variants (SV) causing congenital defects through long-range pathomechanisms ^79^. For example, deletions spanning TAD boundaries can lead to pathological gains in gene expression through enhancer adoption mechanisms whereby enhancers can induce the expression of non-target genes ^79,80^. Our data indicates that the pathological effects of this type of SVs might change (i.e. stronger or weaker induction of non-target genes) depending on whether the deletions include not only CTCF sites, but also nearby developmental genes.

## Acknowledgements

We would like to thank all the Rada-Iglesias lab members for insightful comments and suggestions.

## Methods

### ESC maintenance and differentiation protocol

E14Tg2a (E14) mouse ESCs were cultured on gelatin-coated plates using Knock-out DMEM (Life Technologies, 10829018) supplemented with 15% FBS (Life Technologies, 10082147), leukemia inhibitory factor (LIF), antifungal and antibiotics (Sigma-Aldrich, A5955), β-mercaptoethanol (ThermoFisher Scientific, 21985023), Glutamax (ThermoFisher Scientific, 35050038) and MEM NEAA (ThermoFisher Scientific, 11140035). Cells were cultured at 37°C with 5% C02.

For the NPC differentiation ^59^, ESCs were plated at 20,000 cells/cm^2^ on geltrex-coated plates (ThermoFisher, A1413302) and grown in N2B27 medium: Advanced Dulbecco’s Modified Eagle Medium F12 (Life Technologies, 21041025) and Neurobasal medium (Life Technologies, 12348017) (1:1), supplemented with 1× N2 (R&D Systems, AR009), 1× B27 (Life Technologies, 12587010), 2 mM L-glutamine (Life Technologies, 25030024) and 0.1 mM 2-mercaptoethanol (Life Technologies, 31350010)). During the six days of differentiation, the N2B27 medium was additionally supplemented with the following components: bFGF (ThermoFisher Scientific, PHG0368) 10 ng/ml from day 0 to day 2, Xav939 (Sigma-Aldrich, X3004- 5MG) 5µM from day 2 to day 6, BSA (ThermoFisher Scientific, 15260037) 1mg/ml at day 0 and 40 µg/mL the remaining days.

### RNA isolation, cDNA synthesis and RT-qPCR

Total RNA was isolated using the NZY Total RNA Isolation kit (NZYTech, MB13402) following the manufacturer’s instructions. RNA was reverse transcribed into cDNA using the NZY First-Strand cDNA Synthesis Kit (NZYTech, MB13402). For each reaction, 1 ug of RNA was incubated with 10 µL of NZYRT 2x Master Mix, 2µL of NZYRT Enzyme Mix and nuclease-free water to a total volume of 20µL at 25°C for 10 minutes, followed by 30 minutes at 50°C. The enzyme was heat inactivated at 85°C for 5 minutes. To digest the remaining RNA, 1µL of NZY RnaseH was added to the reaction and incubated at 37°C for 20 minutes.

RT-qPCRs were performed using the CFX 384 detection system (Bio-Rad) using NZYSpeedy qPCR Green Master Mix (2x) (NZYtech, MB224). For each sample, RT-qPCRs were performed as technical triplicates using the primers listed in Data S3. For each cell line, the number of clonal lines and biological replicates analysed in each case is indicated in the corresponding figure legends. For the investigated genes (i.e. Gbx2, Asb18, Six3 and Six2), gene expression fold-changes between each cell line and the corresponding WT cells were calculated using the 2^4-ΔΔCT method, using *Eef1a1* and *Hprt1* as housekeeping genes. Fold-changes are shown as Log2 values.

### CRISPR-Cas9 genome editing

The design of the CRISPR/Cas9 guide sequences (gRNA) was performed using the CRISPR Benchling software tool (https://www.benchling.com/crispr/) (Data S3). In order to increase the cutting efficiency of the gRNA, a guanine nucleotide was added at the first position of the sequence and a restriction site for the BbsI enzyme (R0539L, NEB) was added to the beginning of the gRNA for cloning purposes. For each sgRNA, two oligonucleotides were synthesized (Sigma), annealed and cloned into a CRISPR-Cas9 expression vector (pX330-hCas9-long-chimeric-grna-g2p; gift from Leo Kurian’s laboratory). The hybridized oligos were cloned into the pX330-hCas9-long-chimeric-grna-g2p using 50 ng of BbsI-digested vectors and 1 µl ligase (Thermo Fisher, EL0013) in a total volume of 20 µl. The ligation reaction was incubated for 1 h at room temperature. For each cell line, ESC were transfected with the corresponding pair of gRNAs-Cas9 expressing vectors using Lipofectamine following manufacturer’s recommendations (Thermo Scientific, L3000001). After 24 hours, transfected cells were selected by treating them with puromycin for 48 hours. Single-cell isolation of surviving cells was performed by serial dilution and seeding in 96-well plates. Next, clones with the desired genetic rearrangements (*i.e.* deletions or inversions) were identified by PCR using the primers listed in Data S3. DNA extraction was performed using Lysis Buffer: 25 mM KCl (SigmaAldrich, 27810.295), 5mM TRIS (Sigma-Aldrich, 0497-5KG) pH8.3, 1.25mM MgCl2 (VWR BDH7899-1), 0.225% IGEPAL (Sigma-Aldrich, I8896-50ML) and 0.225% Tween20 (VWR). Proteinase K (ThermoFisher Scientific, EO0492) was added to a final concentration of 0.4 µg/ul before use.

### ChIP-Seq

∼4 × 10^7^ cells were used for RAD21 (abcam, ab154769) and ∼1 × 10^7^ cells for H3K27ac (Active Motif, 39133). Cells were crosslinked with 1% formaldehyde for 10 min at room temperature (RT) and quenched with 0.125 M glycine for 10 min. Cells were consecutively incubated with three different lysis buffers (Buffer 1: 50 mM HEPES, 140 mM NaCl, 1 mM EDTA, 10% glycerol, 0.5% NP-40, 0.25% TX-100; Buffer 2: 10 mM Tris, 200 mM NaCl, 1 mM EDTA, 0.5 mM EGTA; Buffer 3: 10 mM Tris, 100 mM NaCl, 1 mM EDTA, 0.5 mM EGTA, 0.1% Na-deoxycholate, 0.5% N-lauroylsarcosine) in order to isolate chromatin. Next, chromatin was sonicated for 8 minutes in Buffer 3 (20 s on, 30 s off, 25% amplitude) using an EpiShear probe sonicator (Active Motif).

The sonicated chromatin was mixed with 3 µg of H3K27ac antibody or 10 µg of Rad21 antibody and incubated overnight at 4°C. Next, 50-100 µl of Dynabeads (Thermo Fisher, 10-002-D) were aliquoted into a microtube for each ChIP reaction. The dynabeads were washed with 1 ml of cold blocking solution (0,5% BSA and 1x PBS). The antibody bound chromatin was added to the washed beads and incubated for four hours at 4°C. Magnetic beads were washed five times with RIPA buffer (50 mM Hepes, 500 mM LiCl, 1 mM EDTA, 1% NP-40 and 0,7% Na-Deoxycholate). The chromatin was eluted and incubated at 65°C overnight to reverse the crosslinking. Next, samples were treated with 0,2 mg/ml Rnase A and 0,2 mg/ml proteinase K. DNA purification was performed using Zymoclean Gel DNA Recovery kit (Zymo Research, D4008).

Regarding the computational processing of the ChIP-seq data, reads were subject to quality control and trimming of low quality regions and/or adapters using *fastqc* (https://www.bioinformatics.babraham.ac.uk/projects/fastqc/), *MultiQC* ^81^ and *trimmomatic* ^82^. Next, reads were mapped to mm10 reference genome with Bowtie2 ^83^. After read mapping, only reads with a mapping quality above 10 were kept and duplicated reads were removed with SAMtools ^84^. Afterwards, bigwig files were generated with *bamCoverage* from *deepTools* ^85^ applying the reads per genome coverage normalization.

### Capture-C

Capture-C experiments were performed as previously described ^86^. 5 × 10^6^ cells were crosslinked with 2% formaldehyde for 10 minutes and quenched with 0.125 M glycine for 10 minutes. Cells were washed with PBS and resuspended in lysis buffer (10 mM Tris pH 8, 10 mM NaCl, 0.2% NP-40 and 1× protease inhibitors) during 20 minutes on ice. Following centrifugation, the pellet was resuspended in 215 µL 1×CutSmart buffer and transferred to a microcentrifuge tube. The resuspended pellet was mixed with 60 µl 10× CutSmart buffer, 393,5 µl water and 9,5 µl 20% (vol/vol) SDS (0.28% final concentration) (Invitrogen, cat. no. AM9820) followed by 1 h incubation at 37°C while shaking on a thermomixer at 500 rpm (intermittent shaking: 30s on / 30s off). Then, 20% vol/vol Triton X 100 was added at a final concentration of 1.67% vol/vol followed by another incubation at 37°C for 1h while shaking. Chromatin was digested by adding 25 µL NlaIII (250 U, R0125L) and incubating at 37 °C for several hours, followed by addition of another 25 µL of NlaIII and incubation at 37°C overnight. The digested chromatin was ligated with 8 µl (240U) of T4 DNA ligase (Life Tech, cat. no. EL0013) for 18 hours at 16°C. Samples were treated with Proteinase K (3U; Thermo Fisher, cat. no. EO0491) and RNase A (7,5 mU; Roche: 1119915), and DNA was purified using the Qiagen kit (28506). After checking the quality of the digestion and subsequent ligation, chromatin was sonicated for 30 cycles (30 s on, 30 s off, 25% amplitude) using an EpiShear probe sonicator (Active Motif) and DNA samples were purified using AMPure XP SPRI beads (Beckman Coulter, cat. no. A63881). Libraries were prepared using the NEBNext Ultra II kit (New England Biolabs, cat. no. E7645S/L). Index primers set 1 and 2 from the NEBNext Multiplex Oligos for Illumina kit (New England Biolabs, E7335S/L E7500S/L) were incorporated using Herculase II Fusion Polymerase Kit (Agilent, cat. no. 600677). The resulting libraries were pooled (six libraries/pool) and a double Hybridization capture using ssDNA probes was performed following a modified version of the Roche HyperCapture streptavidin pull-down protocol (Roche, cat. no. 09075763001) described in ^86^. Libraries were sequenced using Novaseq6000_150PE_2,25Gb/lib (15 Mreads/lib). For each of the investigated cell lines, Capture-C experiments were performed as two independent biological replicates.

*Capsequm2* (http://apps.molbiol.ox.ac.uk/CaptureC/cgi-bin/CapSequm.cgi) was used to design the ssDNA probes (120 bp probe length). The sequences of the ssDNA probes are listed in Data S3.

### Analysis and quantification of Capture-C data

Capture C reads were subject to quality control and trimming of low quality regions and/or adapters using *fastqc* (https://www.bioinformatics.babraham.ac.uk/projects/fastqc/), *MultiQC* ^81^ and *trimmomatic* ^82^. Next, the reads were processed with capC-MAP ^87^, considering the restriction enzyme *NlaIII* (cutting site CATG), the mm10 reference genome and *normalize = TRUE*. The coordinates (mm10) of the viewpoints (i.e. «targets» according to *capC-MAP* terminology) were:

*Gbx2* SE : chr1 89869929 89870764

*Six3* SE : chr17 85484699 85485038

*Asb18* TSS: chr1 90013014 90013383

*Six2* TSS: chr17 85688402 85689017

After running *capC-MAP*, the normalized pileup bedgraph files for intra-chromosomal contacts for each viewpoint were collected. According to *capC-MAP* documentation, the number of piled-up interactions per restriction fragment are normalized to reads per million, so that the sum of the number of reads associated to each viewpoint genome-wide is equal to one million. Subsequently, for the restriction sites without any detected interactions, a signal equal to 0 was assigned. In addition, only data from the regions *chr1 89228752 90664659* (*Gbx2-Asb18* locus) and *chr17 84890284 86289254* (*Six3-Six2* locus) was considered. Next, the bedgraph files of both replicates were averaged and the resulting bedgraph was converted to a bigwig with the usage of the *bedGraphToBigWig* UCSC tool ^88^. Bigwig subtraction tracks were generated with the usage of *bigwigCompare* from *deepTools* ^85^.

### Annotation of developmental and housekeeping genes

Developmental genes were annotated based on the presence of broad H3K27me3/polycomb domains around their TSS ^89,90^ using a previously described strategy ^59,91^. Briefly, H3K27me3 Chip-Seq fastq files from hESC (GSE24447; H3K27me3: SRR067372, input: SRR067371) and mESC (GSE89209; H3K27me3:SRR4453259, input: SRR4453262) were downloaded. Then, reads were mapped against mm10 and hg19 genomes with bowtie2 and peaks were called with MACS considering the broad peak calling mode ^92^. After peak calling, only peaks with a fold enrichment >3 and q value <0.1 were kept. Next, peaks within 1 Kb of each other were merged using *bedtools*, and associated with a protein coding gene when overlapping a TSS. Subsequently, the knee of the peak size distribution was evaluated with *findiplist()* (inflection R package; https://cran.r-project.org/web/packages/inflection/vignettes/inflection.html). Upon curvature analyses, genes associated with a H3K27me3 peak>7Kb were defined as developmental genes (Mice: n=967; Humans: n=1045).

For the annotation of housekeeping genes, one list for humans and one for mice (Mice: n=3277; Humans: n=2176) were obtained from the *Housekeeping and Reference Transcript Atlas* database ^93^.

### *In silico* analysis of TAD gene density

TAD maps previously generated in different mouse and human cell types were considered. Regarding human TAD maps, the 37 hg19 TAD maps available in the 3D Genome browser were used ^94^. With respect to mice, 14 TAD maps were used: (i) eight mm10 TAD maps available in the 3D Genome browser ^94^, (ii) three mm9 TAD maps (CH12_Lieberman-raw_TADs.txt, cortex_Dixon2012-raw_TADs.txt and mESC_Dixon2012-raw_TADs.txt), also available in 3D genome browser and that were liftover to mm10 using the UCSC liftover tool ^95^, and (iii) three mm10 TAD maps from mESCs, neural progenitors and cortical neurons provided in ^72^. For the eight mm10 TAD maps from the 3D Genome browser, with the exception of the files G1E- ER4.rep1-raw.domains and G1E-ER4.rep2-raw.domains, the prefix “chr” was added to the chromosome name, chr23 was relabeled to chrX and chr24 to chrY.

For each TAD, gene density was computed as the number of TSSs (based on hg19 and mm10 RefSeq curated annotations downloaded through the UCSC Table browser ^96^) located within each TAD divided by the length of the TAD. Subsequently, for each TAD map, the TADs were sorted based on increasing gene densities, and three groups of TADs of equal size were considered: low density (LD), medium density (MD) and high density (HD) TADs. Next, for each TAD map, we computed whether developmental genes were over represented among the genes found within LD TADs using a Fisher test. Lastly, Gene Ontology functional enrichment analyses were performed for two different TAD maps (*hESC Dixon* and *mESC Dixon*, available in 3D genome browser) using the *WebGestalt* R package ^97^ and considering the genes located in the three different groups of TADs (LD, MD and HD) described above. The WebGestalt functional enrichment analysis were performed using the “ORA” (over representation analyses) method and all genes as the reference gene list (group of genes used to compare and compute enrichments).

### In silico analysis of gene distribution within TADs

When computing gene distribution within TADs, the TSSs of genes were taken as a reference. TSS coordinates were obtained from RefSeq curated annotation (downloaded through the UCSC Table browser ^96^) for both human and mice. Gene distribution within TADs was computed in multiple TAD maps independently (37 TAD maps in humans and 14 in mice, see previous methods section).

Each TAD was divided in ten bins of equal sizes. Therefore, the size of each bin is 10% of the size of its corresponding TAD. Regarding the labeling of the bins, when moving from the boundaries of the TAD (TAD start or end coordinates) towards its interior, the first bin was labeled as bin 1, the next one as bin 2 and so on, until reaching bin 5, which is the closest to the center of the TAD (Fig. 1B). The genomic regions located outside TADs (inter-TAD) were assigned to bin 0 (Fig. 1B). For each of the considered TAD maps, the TSSs of genes were assigned to bins attending to their location in the genome.

### Global analysis of insulation strength and CTCF binding at TAD boundaries

The CTCF data used in these analyses were obtained from mESC (GSE36027, CTCF replicates: SRR489719 and SRR489720, input replicates: SRR489731 and SRR489732) and hESCs (GSE116862, CTCF replicates: SRR7506641 and SRR7506642, input replicates: SRR7506652 and SRR7506653). After downloading the corresponding fastq files, reads from both replicates were merged and subject to quality control and trimming of low quality regions and/or adapters using *fastqc*, *MultiQC* ^81^ and *trimmomatic*^82^. Next, reads were mapped to either mm10 or hg19 genomes with *Bowtie2*^83^. After read mapping, only reads with a mapping quality above 10 were kept and duplicated reads were removed with *SAMtools* ^84^. Afterwards, CTCF bigwig files were generated with *bamCoverage* from *deepTools* applying the reads per genome coverage normalization ^85^. CTCF peaks were called with MACS2 ^92^ and only those peaks with a fold change > 4 and q value < 0.01 were considered. Lastly, using the coordinates of the CTCF peaks and the CTCF bigwig files, the maximum intensity of each CTCF peak was calculated using the *bigWigAverageOverBed* UCSC binary tool.

The insulation score and boundary strength datasets used for our analyses were obtained from different sources and through various procedures. Regarding mESC data, a bigwig file (file ID: 4DNFIMVJ2YV3) with mm10 insulation scores obtained from previously generated Hi-C data ^72^ was downloaded from the 4D Nucleome data portal ^98^. In addition, a bed file (file id: 4DNFI1S7FI1U) with boundary strength values as computed by *cooltools* ^99^ for the bins mapped as TAD boundaries was also downloaded from the 4D Nucleome data portal ^98^. Regarding hESC data, *.hic* files from two replicates (GSM3262956 and GSM3262957 ^23^) were downloaded from GEO Next, *.hic* files were converted to *.cool* format with the *hic2cool* software (https://github.com/4dn-dcic/hic2cool) considering the 10 Kb contact resolution matrix. Afterwards both replicates were merged with *cooler merge* and normalised with *cooler balance* ^100^. Subsequently, insulation scores were computed with *cooltools* (based on the diamond insulation score technique) using a window size of 100 Kb. In addition a bigwig file was also created in this step with *cooltools* by specifying the *–bigwig* option. Moreover, boundary strength metrics were also computed by *cooltools* for the bins considered as TAD boundaries. In order to make these boundary strength values more comparable to those obtained from the 4D Nucleome data portal, only TAD boundaries with a strength larger than 0.2 were considered, while a strength value=0 was assigned to the remaining boundaries

Once ready, the insulation score, boundary strength and CTCF datasets were used to compute several metrics for TAD boundaries in both mice and humans. TAD boundaries were defined using the start and end coordinates of TADs previously identified in mESC ^72^ and hESC ^101^. Next, each TAD boundary was expanded by +/- 50 Kb and the following metrics were calculated within the resulting 100 Kb window: (i) the number of overlapping CTCF peaks, (ii) the CTCF aggregated signal (sum of the CTCF bigwig signal for all the peaks overlapping with the 100kb window), (iii) the insulation score (minimum insulation score value in the 100kb window computed with *bigWigAverageOverBed*) and (iv) the boundary strength (maximum boundary strength value computed by *cooltools* for any bin in the 100kb window). In addition, TAD boundaries were classified as “Developmental” or “Other”: “Developmental” boundaries were defined as those associated with a developmental gene located in bin 1 (Fig.1 1B); “Other” included all the remaining boundaries. Moreover, a set of 5000 “random” TAD boundaries was generated by randomly selecting 5000 regions in the mouse and human genomes.

### *In silico* analysis of the distribution of CTCF sites around genes located close to TAD boundaries

Gene located close to TAD boundaries were defined as those assigned to bin 1 (Fig. 1B) according to TAD maps previously generated in mESC ^72^ and hESC ^101^. Then, for each “bin 1” gene, the number of CTCF peaks (obtained as described in the previous section) located within a +/- 100 Kb window around its TSS was calculated. The 100 Kb window extending from the gene TSS towards the TAD boundary was defined as the “outer window”, while the 100 Kb window extending from the gene TSS towards the center of the TAD was defined as the “inner window” (Fig. 2C). After calculating the number of CTCF peaks in the “inner” and “outer” windows, we computed the *ΔCTCFpeaks@Bdry* metric for each gene as the number of CTCF peaks located in the “inner window” minus the number of CTCF peaks located in the “outer window”. In addition, the CTCF bigwig signals associated with the CTCF peaks were used to calculate the *ΔCTCFsignal@Bdry* metric for each gene as the aggregated signal for all the CTCF peaks located in the “inner window” minus the aggregated signal for all the CTCF peaks located in the “outer window”.

## References

1. Furlong, E. E. M. & Levine, M. Developmental enhancers and chromosome topology. Science 361, 1341–1345 (2018).

2. Lagha, M., Bothma, J. P. & Levine, M. Mechanisms of transcriptional precision in animal development. Trends Genet. 28, 409–416 (2012).

3. Long, H. K., Prescott, S. L. & Wysocka, J. Ever-Changing Landscapes: Transcriptional Enhancers in Development and Evolution. Cell 167, 1170– 1187 (2016).

4. West, A. G., Gaszner, M. & Felsenfeld, G. Insulators: many functions, many mechanisms. Genes Dev. 16, 271–288 (2002).

5. Hnisz, D., Day, D. S. & Young, R. A. Insulated Neighborhoods: Structural and Functional Units of Mammalian Gene Control. Cell 167, 1188–1200 (2016).

6. Bell, A. C., West, A. G. & Felsenfeld, G. The Protein CTCF Is Required for the Enhancer Blocking Activity of Vertebrate Insulators. Cell 98, 387–396 (1999).

7. Geyer, P. K. & Corces, V. G. DNA position-specific repression of transcription by a Drosophila zinc finger protein. Genes Dev. 6, 1865–1873 (1992).

8. Cavalheiro, G. R. et al. CTCF, BEAF-32, and CP190 are not required for the establishment of TADs in early *Drosophila* embryos but have locus-specific roles. Sci. Adv. 9, eade1085 (2023).

9. Chathoth, K. T. et al. The role of insulators and transcription in 3D chromatin organization of flies. Genome Res. 32, 682–698 (2022).

10. Kaushal, A. et al. Essential role of Cp190 in physical and regulatory boundary formation. Sci. Adv. 8, eabl8834 (2022).

11. Nora, E. P. et al. Targeted Degradation of CTCF Decouples Local Insulation of Chromosome Domains from Genomic Compartmentalization. Cell 169, 930–944.e22 (2017).

12. Ohlsson, R., Renkawitz, R. & Lobanenkov, V. CTCF is a uniquely versatile transcription regulator linked to epigenetics and disease. Trends Genet. 17, 520–527 (2001).

13. Rowley, M. J. et al. Evolutionarily Conserved Principles Predict 3D Chromatin Organization. Mol. Cell 67, 837–852.e7 (2017).

14. Dixon, J. R. et al. Topological domains in mammalian genomes identified by analysis of chromatin interactions. Nature 485, 376–380 (2012).

15. Harmston, N. et al. Topologically associating domains are ancient features that coincide with Metazoan clusters of extreme noncoding conservation. Nat. Commun. 8, 441 (2017).

16. Rao, S. S. P. et al. Cohesin Loss Eliminates All Loop Domains. Cell 171, 305–320.e24 (2017).

17. Schwarzer, W. et al. Two independent modes of chromatin organization revealed by cohesin removal. Nature 551, 51–56 (2017).

18. Kaushal, A. et al. CTCF loss has limited effects on global genome architecture in Drosophila despite critical regulatory functions. Nat. Commun. 12, 1011 (2021).

19. Ramírez, F. et al. High-resolution TADs reveal DNA sequences underlying genome organization in flies. Nat. Commun. 9, 189 (2018).

20. Banigan, E. J. et al. Transcription shapes 3D chromatin organization by interacting with loop extrusion. Proc. Natl. Acad. Sci. U. S. A. 120, e2210480120 (2023).

21. Bozhilov, Y. K. et al. A gain-of-function single nucleotide variant creates a new promoter which acts as an orientation-dependent enhancer-blocker. Nat. Commun. 12, 3806 (2021).

22. Zhang, S. et al. RNA polymerase II is required for spatial chromatin reorganization following exit from mitosis. Sci. Adv. 7, eabg8205 (2021).

23. Zhang, Y. et al. Transcriptionally active HERV-H retrotransposons demarcate topologically associating domains in human pluripotent stem cells. Nat. Genet. 51, 1380–1388 (2019).

24. Zhang, D. et al. Alteration of genome folding via contact domain boundary insertion. Nat. Genet. 52, 1076–1087 (2020).

25. Beccari, L. et al. Dbx2 regulation in limbs suggests interTAD sharing of enhancers. Dev. Dyn. Off. Publ. Am. Assoc. Anat. 250, 1280–1299 (2021).

26. Luppino, J. M. et al. Cohesin promotes stochastic domain intermingling to ensure proper regulation of boundary-proximal genes. Nat. Genet. 52, 840– 848 (2020).

27. Hung, T.-C., Kingsley, D. M. & Boettiger, A. Boundary stacking interactions enable cross-TAD enhancer-promoter communication during limb development. http://biorxiv.org/lookup/doi/10.1101/2023.02.06.527380(2023) doi:10.1101/2023.02.06.527380.

28. Chakraborty, S. et al. Enhancer–promoter interactions can bypass CTCF-mediated boundaries and contribute to phenotypic robustness. Nat. Genet. 55, 280–290 (2023).

29. Balasubramanian, D. et al. Enhancer–promoter interactions can form independently of genomic distance and be functional across TAD boundaries. Nucleic Acids Res. gkad1183 (2023) doi:10.1093/nar/gkad1183.

30. Chang, L.-H., Ghosh, S. & Noordermeer, D. TADs and Their Borders: Free Movement or Building a Wall? J. Mol. Biol. 432, 643–652 (2020).

31. Chang, L.-H. et al. Multi-feature clustering of CTCF binding creates robustness for loop extrusion blocking and Topologically Associating Domain boundaries. Nat. Commun. 14, 5615 (2023).

32. Bartman, C. R., Hsu, S. C., Hsiung, C. C.-S., Raj, A. & Blobel, G. A. Enhancer Regulation of Transcriptional Bursting Parameters Revealed by Forced Chromatin Looping. Mol. Cell 62, 237–247 (2016).

33. Lower, K. M. et al. Adventitious changes in long-range gene expression caused by polymorphic structural variation and promoter competition. Proc. Natl. Acad. Sci. U. S. A. 106, 21771–21776 (2009).

34. Topfer, S. K. et al. Disrupting the adult globin promoter alleviates promoter competition and reactivates fetal globin gene expression. Blood 139, 2107– 2118 (2022).

35. Ohtsuki, S., Levine, M. & Cai, H. N. Different core promoters possess distinct regulatory activities in the Drosophila embryo. Genes Dev. 12, 547–556 (1998).

36. Ohtsuki, S. & Levine, M. GAGA mediates the enhancer blocking activity of the *eve* promoter in the *Drosophila* embryo. Genes Dev. 12, 3325–3330 (1998).

37. Cai, H. N., Zhang, Z., Adams, J. R. & Shen, P. Genomic context modulates insulator activity through promoter competition. Development 128, 4339– 4347 (2001).

38. Palstra, R., De Laat, W. & Grosveld, F. Chapter 4 β-Globin Regulation and Long-Range Interactions. in Advances in Genetics vol. 61 107–142 (Elsevier, 2008).

39. Oudelaar, A. M. et al. A revised model for promoter competition based on multi-way chromatin interactions at the α-globin locus. Nat. Commun. 10, 5412 (2019).

40. Fukaya, T., Lim, B. & Levine, M. Enhancer Control of Transcriptional Bursting. Cell 166, 358–368 (2016).

41. Lim, B. & Levine, M. S. Enhancer-promoter communication: hubs or loops? Curr. Opin. Genet. Dev. 67, 5–9 (2021).

42. Deng, H. & Lim, B. Shared Transcriptional Machinery at Homologous Alleles Leads to Reduced Transcription in Early Drosophila Embryos. Front. Cell Dev. Biol. 10, 912838 (2022).

43. Sabi, R. & Tuller, T. Modelling and measuring intracellular competition for finite resources during gene expression. J. R. Soc. Interface 16, 20180887 (2019).

44. Cruz-Molina, S. et al. PRC2 Facilitates the Regulatory Topology Required for Poised Enhancer Function during Pluripotent Stem Cell Differentiation. Cell Stem Cell 20, 689–705.e9 (2017).

45. Laugsch, M. et al. Modeling the Pathological Long-Range Regulatory Effects of Human Structural Variation with Patient-Specific hiPSCs. Cell Stem Cell 24, 736–752.e12 (2019).

46. Yan, K.-K., Lou, S. & Gerstein, M. MrTADFinder: A network modularity based approach to identify topologically associating domains in multiple resolutions. PLoS Comput. Biol. 13, e1005647 (2017).

47. Pachano, T., Haro, E. & Rada-Iglesias, A. Enhancer-gene specificity in development and disease. Development 149, dev186536 (2022).

48. Lenhard, B., Sandelin, A. & Carninci, P. Metazoan promoters: emerging characteristics and insights into transcriptional regulation. Nat. Rev. Genet. 13, 233–245 (2012).

49. Simeone, A. Positioning the isthmic organizer where Otx2 and Gbx2meet. Trends Genet. TIG 16, 237–240 (2000).

50. Tai, C.-I. & Ying, Q.-L. Gbx2, a LIF/Stat3 target, promotes reprogramming to and retention of the pluripotent ground state. J. Cell Sci. 126, 1093–1098 (2013).

51. Kubo, N. et al. Promoter-proximal CTCF binding promotes distal enhancer-dependent gene activation. Nat. Struct. Mol. Biol. 28, 152–161 (2021).

52. Anania, C. et al. In vivo dissection of a clustered-CTCF domain boundary reveals developmental principles of regulatory insulation. Nat. Genet. 54, 1026–1036 (2022).

53. Huang, H. et al. CTCF mediates dosage- and sequence-context-dependent transcriptional insulation by forming local chromatin domains. Nat. Genet. 53, 1064–1074 (2021).

54. Karr, J. P., Ferrie, J. J., Tjian, R. & Darzacq, X. The transcription factor activity gradient (TAG) model: contemplating a contact-independent mechanism for enhancer-promoter communication. Genes Dev. 36, 7–16 (2022).

55. Zuin, J. et al. Nonlinear control of transcription through enhancer-promoter interactions. Nature 604, 571–577 (2022).

56. Kane, L. et al. Cohesin is required for long-range enhancer action at the Shh locus. Nat. Struct. Mol. Biol. 29, 891–897 (2022).

57. Gómez-Marín, C. et al. Evolutionary comparison reveals that diverging CTCF sites are signatures of ancestral topological associating domains borders. Proc. Natl. Acad. Sci. 112, 7542–7547 (2015).

58. O’Brien, L. L. et al. Transcriptional regulatory control of mammalian nephron progenitors revealed by multi-factor cistromic analysis and genetic studies. PLOS Genet. 14, e1007181 (2018).

59. Pachano, T. et al. Orphan CpG islands amplify poised enhancer regulatory activity and determine target gene responsiveness. Nat. Genet. 53, 1036– 1049 (2021).

60. Batut, P. J. et al. Genome organization controls transcriptional dynamics during development. Science 375, 566–570 (2022).

61. Elango, N. & Yi, S. V. Functional relevance of CpG island length for regulation of gene expression. Genetics 187, 1077–1083 (2011).

62. Ngan, C. Y. et al. Chromatin interaction analyses elucidate the roles of PRC2- bound silencers in mouse development. Nat. Genet. 52, 264–272 (2020).

63. de Laat, W. & Duboule, D. Topology of mammalian developmental enhancers and their regulatory landscapes. Nature 502, 499–506 (2013).

64. Whyte, W. A. et al. Master transcription factors and mediator establish super-enhancers at key cell identity genes. Cell 153, 307–319 (2013).

65. Ringel, A. R. et al. Repression and 3D-restructuring resolves regulatory conflicts in evolutionarily rearranged genomes. Cell 185, 3689–3704.e21 (2022).

66. Roider, H. G., Lenhard, B., Kanhere, A., Haas, S. A. & Vingron, M. CpG-depleted promoters harbor tissue-specific transcription factor binding signals—implications for motif overrepresentation analyses. Nucleic Acids Res. 37, 6305–6315 (2009).

67. Dejosez, M. et al. Regulatory architecture of housekeeping genes is driven by promoter assemblies. Cell Rep. 42, 112505 (2023).

68. Kikuta, H. et al. Genomic regulatory blocks encompass multiple neighboring genes and maintain conserved synteny in vertebrates. Genome Res. 17, 545– 555 (2007).

69. Wu, H.-J. et al. Topological isolation of developmental regulators in mammalian genomes. Nat. Commun. 12, 4897 (2021).

70. Rinzema, N. J. et al. Building regulatory landscapes reveals that an enhancer can recruit cohesin to create contact domains, engage CTCF sites and activate distant genes. Nat. Struct. Mol. Biol. 29, 563–574 (2022).

71. Bateman, J. R. & Johnson, J. E. Altering enhancer–promoter linear distance impacts promoter competition in *cis* and in *trans*. Genetics 222, iyac098 (2022).

72. Bonev, B. et al. Multiscale 3D Genome Rewiring during Mouse Neural Development. Cell 171, 557–572.e24 (2017).

73. Waymack, R., Gad, M. & Wunderlich, Z. Molecular competition can shape enhancer activity in the Drosophila embryo. iScience 24, 103034 (2021).

74. Xiao, J. Y., Hafner, A. & Boettiger, A. N. How subtle changes in 3D structure can create large changes in transcription. eLife 10, e64320 (2021).

75. Paliou, C. et al. Preformed chromatin topology assists transcriptional robustness of *Shh* during limb development. Proc. Natl. Acad. Sci. 116, 12390–12399 (2019).

76. Chen, L.-F. et al. Structural elements promote architectural stripe formation and facilitate ultra-long-range gene regulation at a human disease locus. Mol. Cell 83, 1446–1461.e6 (2023).

77. Andrey, G. et al. A switch between topological domains underlies HoxD genes collinearity in mouse limbs. Science 340, 1234167 (2013).

78. Hafner, A. et al. Loop stacking organizes genome folding from TADs to chromosomes. Mol. Cell 83, 1377–1392.e6 (2023).

79. Spielmann, M., Lupiáñez, D. G. & Mundlos, S. Structural variation in the 3D genome. Nat. Rev. Genet. 19, 453–467 (2018).

80. Lupiáñez, D. G. et al. Disruptions of topological chromatin domains cause pathogenic rewiring of gene-enhancer interactions. Cell 161, 1012–1025 (2015).

81. Ewels, P., Magnusson, M., Lundin, S. & Käller, M. MultiQC: summarize analysis results for multiple tools and samples in a single report. Bioinforma. Oxf. Engl. 32, 3047–3048 (2016).

82. Bolger, A. M., Lohse, M. & Usadel, B. Trimmomatic: a flexible trimmer for Illumina sequence data. Bioinforma. Oxf. Engl. 30, 2114–2120 (2014).

83. Langmead, B. & Salzberg, S. L. Fast gapped-read alignment with Bowtie 2. Nat. Methods 9, 357–359 (2012).

84. Li, H. et al. The Sequence Alignment/Map format and SAMtools. Bioinforma. Oxf. Engl. 25, 2078–2079 (2009).

85. Ramírez, F. et al. deepTools2: a next generation web server for deep-sequencing data analysis. Nucleic Acids Res. 44, W160–165 (2016).

86. Downes, D. J. et al. Capture-C: a modular and flexible approach for high-resolution chromosome conformation capture. Nat. Protoc. 17, 445–475 (2022).

87. Buckle, A., Gilbert, N., Marenduzzo, D. & Brackley, C. A. capC-MAP: software for analysis of Capture-C data. Bioinforma. Oxf. Engl. 35, 4773–4775 (2019).

88. Kent, W. J., Zweig, A. S., Barber, G., Hinrichs, A. S. & Karolchik, D. BigWig and BigBed: enabling browsing of large distributed datasets. Bioinforma. Oxf. Engl. 26, 2204–2207 (2010).

89. Rehimi, R. et al. Epigenomics-Based Identification of Major Cell Identity Regulators within Heterogeneous Cell Populations. Cell Rep. 17, 3062–3076 (2016).

90. Shim, W. J. et al. Conserved Epigenetic Regulatory Logic Infers Genes Governing Cell Identity. Cell Syst. 11, 625–639.e13 (2020).

91. Sánchez-Gaya, V. & Rada-Iglesias, A. POSTRE: a tool to predict the pathological effects of human structural variants. Nucleic Acids Res. 51, e54 (2023).

92. Feng, J., Liu, T., Qin, B., Zhang, Y. & Liu, X. S. Identifying ChIP-seq enrichment using MACS. Nat. Protoc. 7, 1728–1740 (2012).

93. Hounkpe, B. W., Chenou, F., de Lima, F. & De Paula, E. V. HRT Atlas v1.0 database: redefining human and mouse housekeeping genes and candidate reference transcripts by mining massive RNA-seq datasets. Nucleic Acids Res. 49, D947–D955 (2021).

94. Wang, Y. et al. The 3D Genome Browser: a web-based browser for visualizing 3D genome organization and long-range chromatin interactions. Genome Biol. 19, 151 (2018).

95. Hinrichs, A. S. et al. The UCSC Genome Browser Database: update 2006. Nucleic Acids Res. 34, D590–598 (2006).

96. Karolchik, D. et al. The UCSC Table Browser data retrieval tool. Nucleic Acids Res. 32, D493–496 (2004).

97. Liao, Y., Wang, J., Jaehnig, E. J., Shi, Z. & Zhang, B. WebGestalt 2019: gene set analysis toolkit with revamped UIs and APIs. Nucleic Acids Res. 47, W199– W205 (2019).

98. Reiff, S. B. et al. The 4D Nucleome Data Portal as a resource for searching and visualizing curated nucleomics data. Nat. Commun. 13, 2365 (2022).

99. Open2C et al. Cooltools: enabling high-resolution Hi-C analysis in Python. http://biorxiv.org/lookup/doi/10.1101/2022.10.31.514564 (2022) doi:10.1101/2022.10.31.514564.

100. Abdennur, N. & Mirny, L. A. Cooler: scalable storage for Hi-C data and other genomically labeled arrays. Bioinforma. Oxf. Engl. 36, 311–316 (2020).

101. Dixon, J. R. et al. Chromatin architecture reorganization during stem cell differentiation. Nature 518, 331–336 (2015).

102. Pope, B. D. et al. Topologically associating domains are stable units of replication-timing regulation. Nature 515, 402–405 (2014).

