## Supplementary Figures for "Synergistic insulation of regulatory domains by developmental genes and clusters of CTCF sites"

***Supplementary Figure 1.*** *Hi-C profiles at representative examples of mouse "developmental" TADs.*

***Supplementary Figure 2.*** *Hi-C profiles at representative examples of human "developmental" TADs.*

***Supplementary Figure 3.*** *Enriched GO categories for human TADs with different gene densities.*

***Supplementary Figure 4.*** *Developmental genes and clusters of CTCF sites are sequentially organized near human TAD boundaries.*

***Supplementary Figure 5.*** *Hi-C profiles around mouse genes previously reported to bypass TAD boundaries.*

***Supplementary Figure 6.*** *Hi-C profiles around human genes previously reported to bypass TAD boundaries.*

***Supplementary Figure 7.*** *Δ3XCTCF, 71Kb INV and Δ3XCTCF:71Kb INV re-arrangements generated at the Gbx2/Asb18 locus.*

***Supplementary Figure 8.*** *Capture-C experiments in Δ3XCTCF, 71Kb INV and Δ3XCTCF:71Kb INV mESC using the Asb18 promoter as a viewpoint.*

***Supplementary Figure 9.*** *Genotyping of the ΔPromGbx2 and Δ3XCTCF:ΔPromGbx2 re-arrangements generated at the Gbx2/Asb18 locus.*

***Supplementary Figure 10.*** *Capture-C experiments in Δ3XCTCF, ΔPromGbx2 and Δ3XCTCF:ΔPromGbx2 mESC using the Asb18 promoter as a viewpoint.*

***Supplementary Figure 11.*** *RAD21 and H3K27ac profiles in ESC with genomic re-arrangements within the Gbx2/Asb18 locus.*

***Supplementary Figure 12.*** *Genotyping of the Δ6XCTCF, 156Kb INV and Δ6XCTCF:156Kb INV re-arrangements generated at the Six3/Six2 locus.*

***Supplementary Figure 13.*** *Genotyping of the 226Kb INV ESC lines.*

***Supplementary Figure 14.*** *Genotyping of the Δ6XCTCF:Six2$^{-/-}$ ESC lines.*

***Supplementary Figure 15.*** *Genotyping of the Six3$^{-/-}$ and Δ6XCTCF:Six3$^{-/-}$ deletions generated at the Six3/Six2 locus.*

***Supplementary Figure 16.*** *Capture-C experiments in Δ6XCTCF, Six3$^{-/-}$ and Δ6XCTCF:Six3$^{-/-}$ NPC using the Six2 promoter as a viewpoint.*

***Supplementary Figure 17.*** *RAD21 and H3K27ac profiles in NPC with genomic re-arrangements within the Six3/Six2 locus.*

***Supplementary Figure 18.*** *Genotyping of the Δ4XCTCF and Δ4XCTCF:Six3$^{-/-}$ deletions generated at the Six3/Six2 locus.*

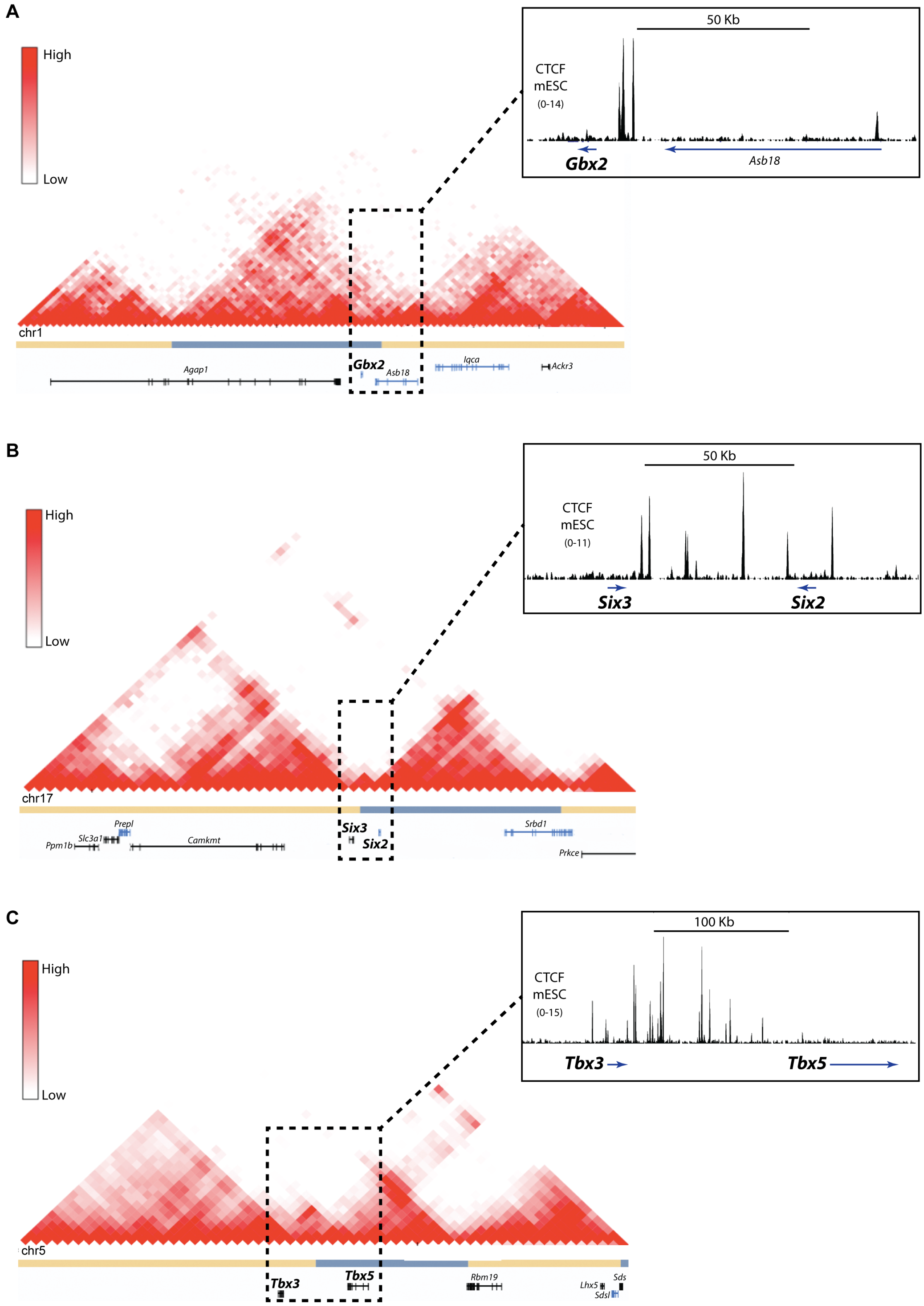


***Supplementary Fig. 1: Hi-C profiles at representative examples of mouse "developmental" TADs. (A-C)*** *Hi-C data from mESC [72] around (A) Gbx2, (B) Six3/Six2 and (C) Tbx3/Tbx5, which serve as representative examples of mouse developmental genes located near TAD boundaries and within gene-poor TADs. CTCF ChIP-seq profiles in mESC [102] are shown around the TAD boundaries located near Gbx2, Six3/Six2 and Tbx3/Tbx5.*

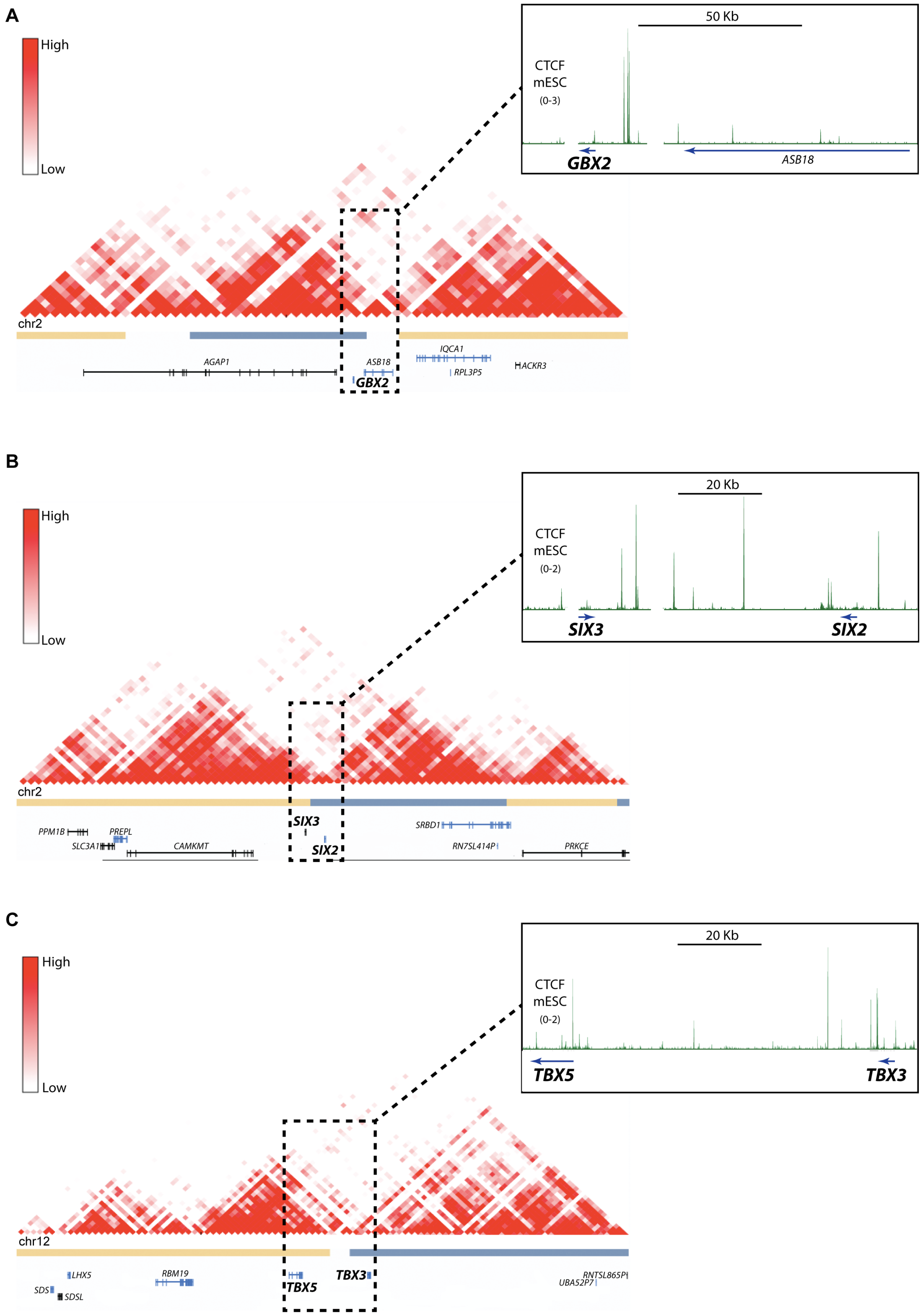


***Supplementary Fig. 2: Hi-C profiles at representative examples of human "developmental" TADs. (A-C)*** *Hi-C data from hESC [101] around (A) GBX2, (B) SIX3/SIX2 and (C) TBX3/TBX5, which serve as representative examples of human developmental genes located near TAD boundaries and within gene-poor TADs. CTCF ChIP-seq profiles in hESC [102] are shown around the TAD boundaries located near GBX2, SIX3/SIX2 and TBX3/TBX5.*

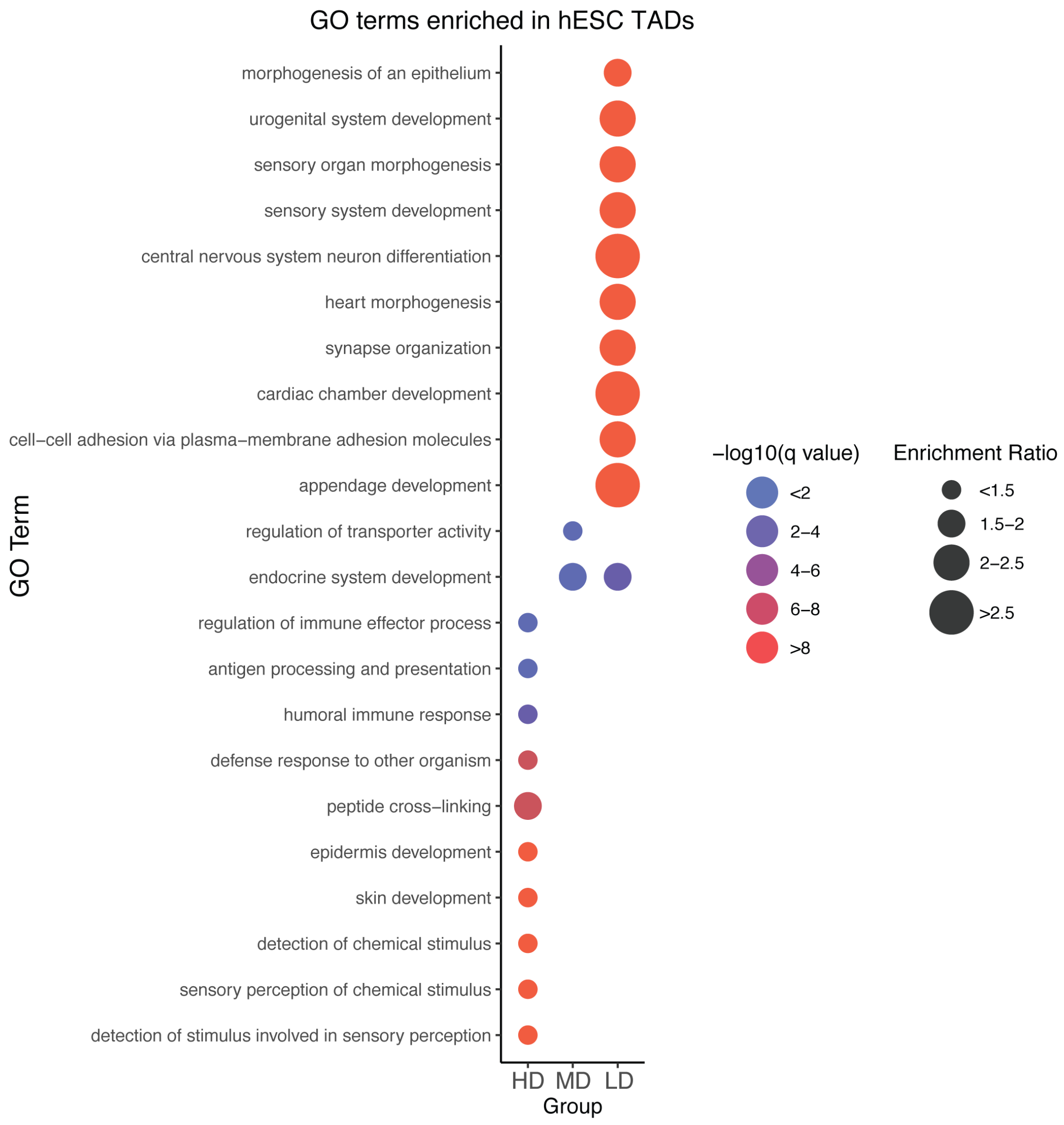


***Supplementary Fig. 3: Enriched GO categories for human TADs with different gene densities.*** *TADs previously identified in hESC [101] were classified in three different groups based on their gene density: High Density (HD), Medium Density (MD) and Low Density (LD). Then, the genes present within each TAD group were subject to GO enrichment analysis. For the LD (n=130 enriched GO terms) group, only the top 10 most significantly enriched GO terms are shown, while for the MD (n=10 enriched GO terms) and HD (n=2 enriched GO terms) groups all the significantly enriched GO terms are presented.*

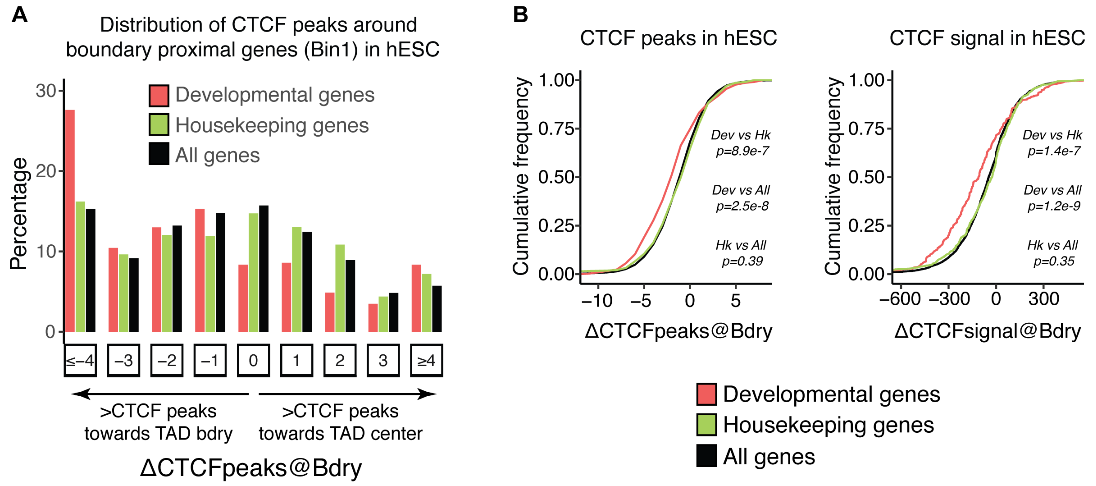


***Supplementary Fig. 4: Developmental genes and clusters of CTCF sites are sequentially organized near human TAD boundaries. (A)*** *Histogram showing the distribution of ΔCTCFpeaks@Bdry values in hESC for different types of genes (developmental, housekeeping, all) located close to TAD boundaries (Bin 1 in Fig 1B).* ***(B)*** *Cumulative distribution plots for ΔCTCFpeaks@Bdry (left) and ΔCTCFsignal@Bdry (right) values in hESC show that for developmental genes the number of CTCF peaks and CTCF signals is significantly more skewed towards negative values than for the other considered gene categories. The ΔCTCFpeaks@Bdry and ΔCTCFsignal@Bdry metrics were calculated in hESC using previously reported TAD maps and CTCF ChIP-seq profiles (see Methods). P-values were calculated using unpaired two-sided Wilcoxon tests with Bonferroni correction for multiple testing.*

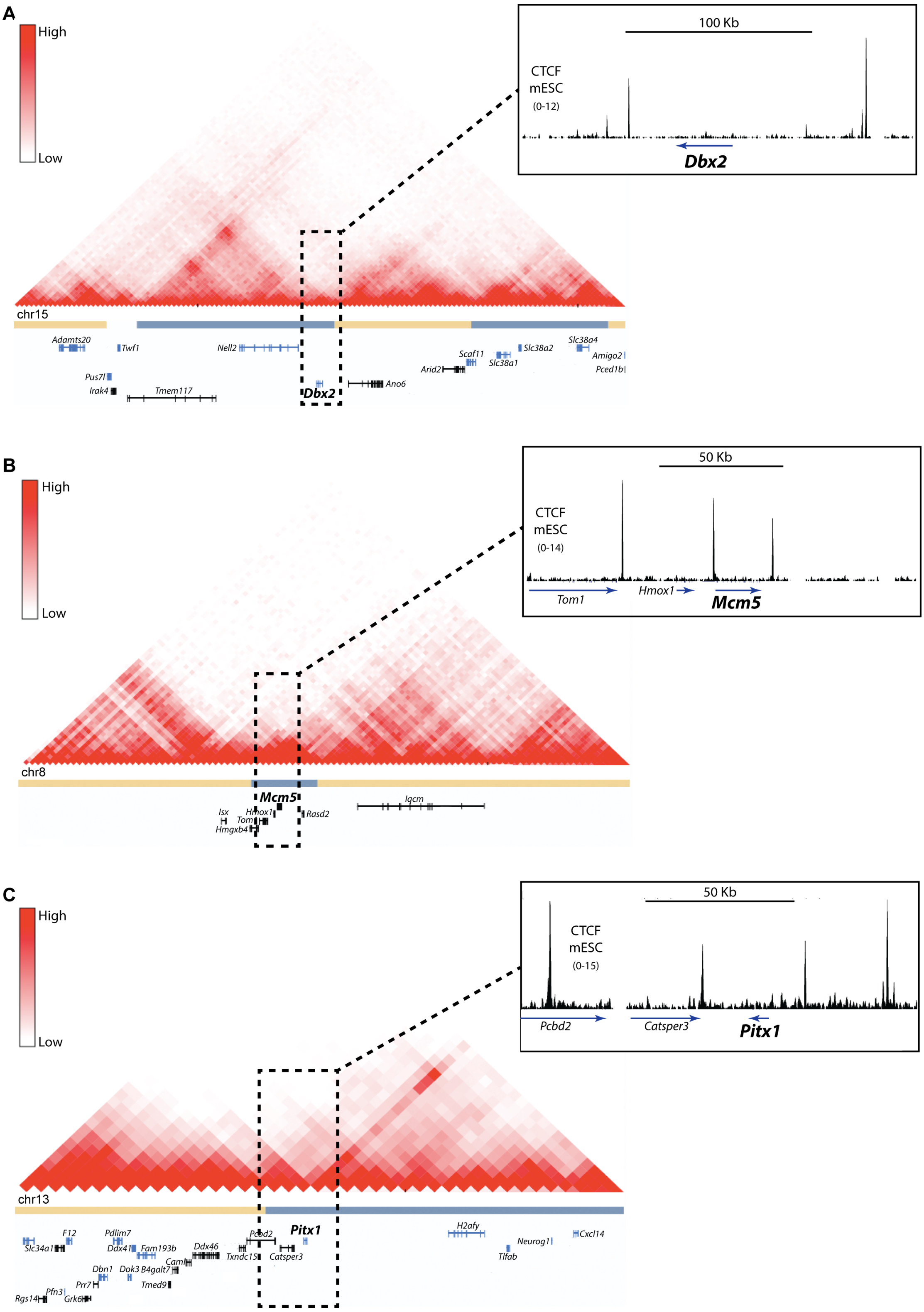


***Supplementary Fig. 5: Hi-C profiles around mouse genes previously reported to bypass TAD boundaries. (A-C)*** *Hi-C data [72] around (A) Dbx2, (B) Mcm5 and (C) Pitx1, which were previously reported to bypass TAD boundaries[25–27]. CTCF ChIP-seq profiles in mESC [102] are shown around the TAD boundaries located near Dbx2, Mcm5 and Pitx1.*

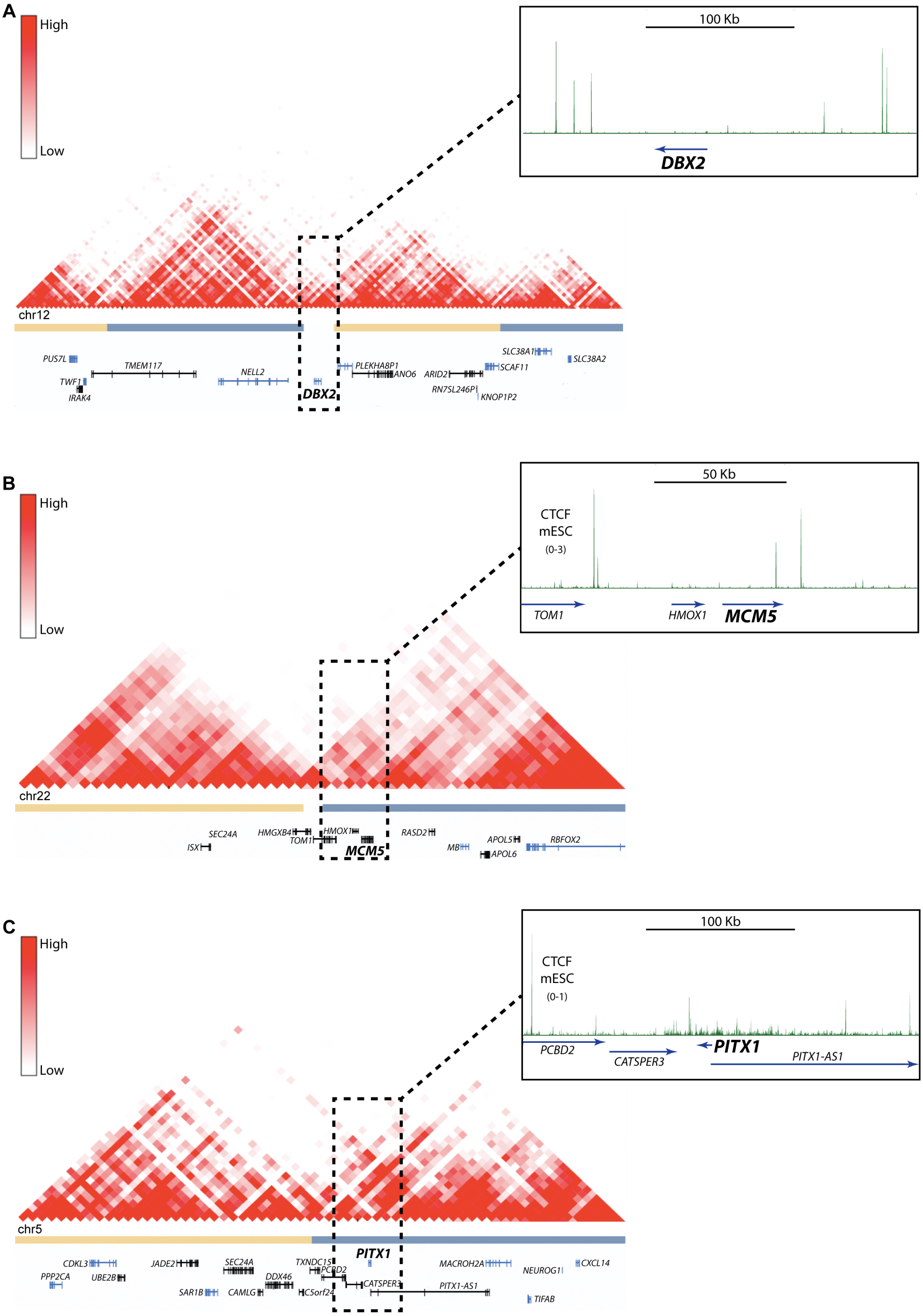


***Supplementary Fig. 6: Hi-C profiles around human genes previously reported to bypass TAD boundaries. (A-C)*** *Hi-C data [101] around (A) DBX2, (B) MCM5 and (C) PITX1, which were previously reported to bypass TAD boundaries[25–27]. CTCF ChIP-seq profiles in hESC [102] are shown around the TAD boundaries located near DBX2, MCM5 and PITX1.*

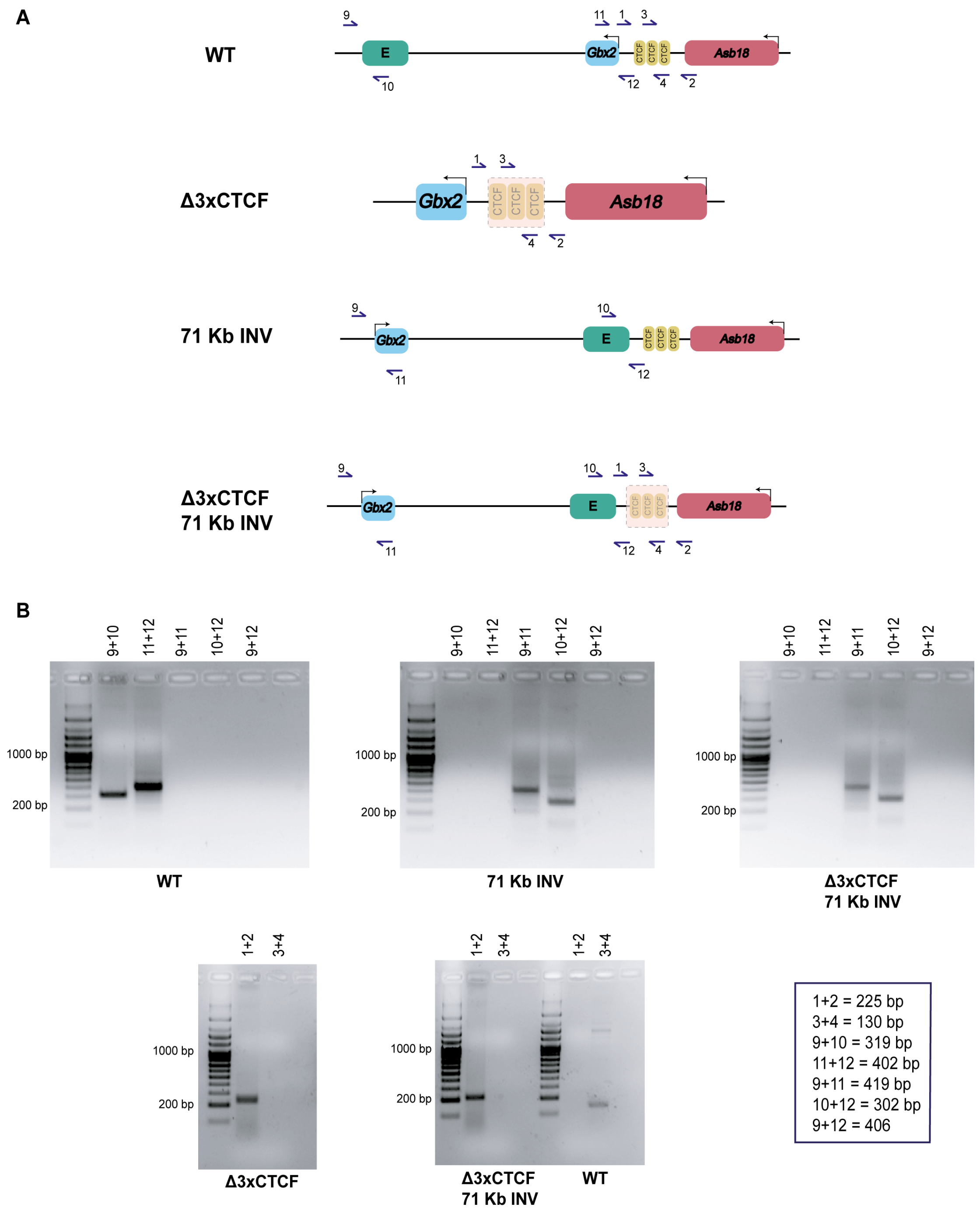


***Supplementary Fig. 7: Genotyping of the Δ3XCTCF, 71Kb INV and Δ3XCTCF:71Kb INV re-arrangements generated at the Gbx2/Asb18 locus. (A)*** *Graphical overview of the PCR-based strategy used to genotype the Δ3XCTCF, 71Kb INV and Δ3XCTCF:71Kb INV genomic re-arrangements described in Fig 3. The horizontal arrows and accompanying numbers represent PCR primers.* ***(B)*** *Representative PCR genotyping results obtained for ESC lines that were either WT or homozygous for the Δ3XCTCF, 71 KB INV and Δ3XCTCF:71Kb INV genomic re-arrangements using the indicated primer pair combinations. The expected sizes of the amplicons obtained with each PCR primer combination are shown at the bottom right corner.*

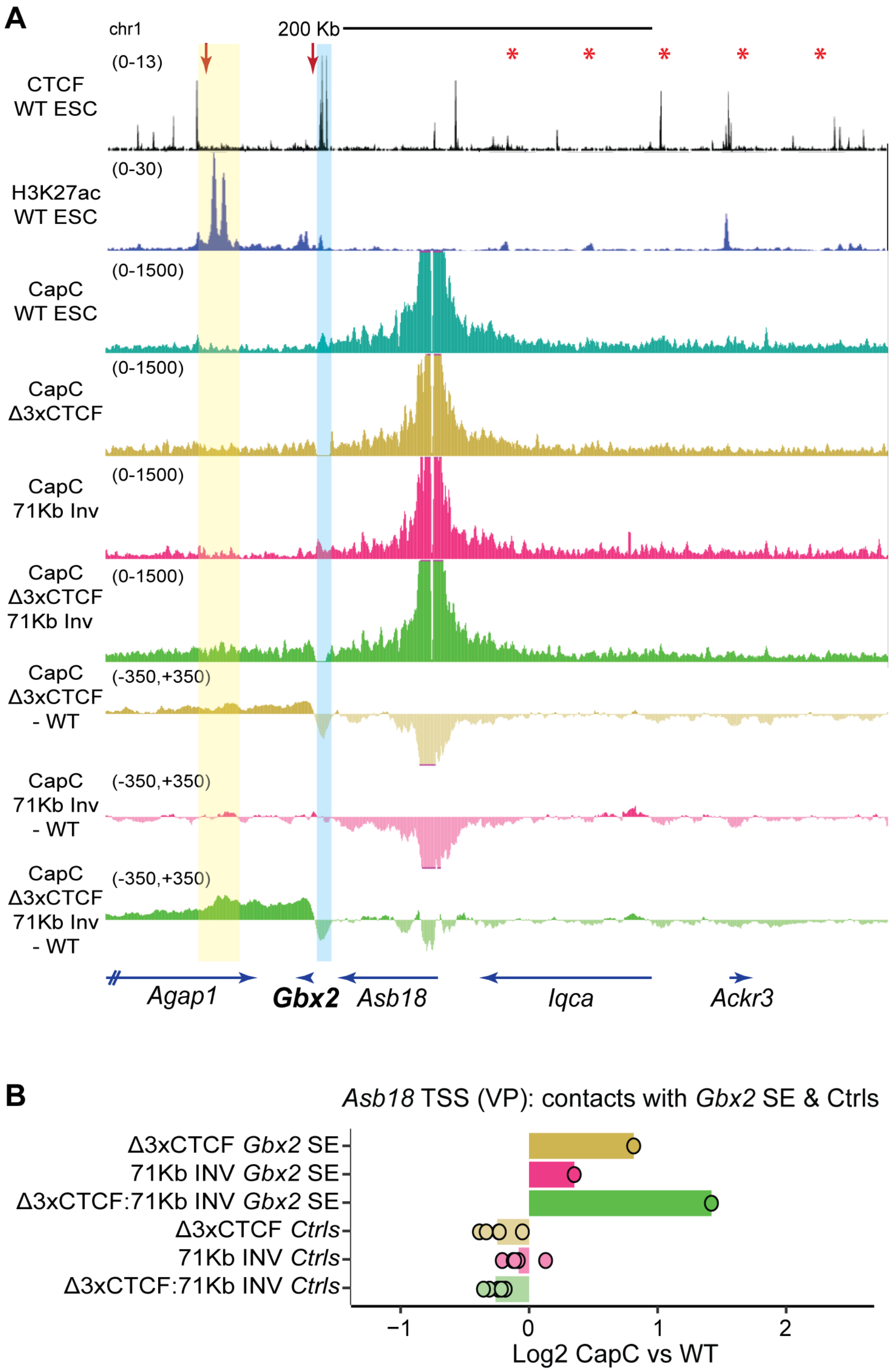


***Supplementary Fig. 8: Capture-C experiments in Δ3XCTCF, 71Kb INV and Δ3XCTCF:71Kb INV mESC using the Asb18 promoter as a viewpoint. (A)*** *Capture-C experiments were performed as two biological replicates in WT, Δ3XCTCF, 71Kb INV and Δ3XCTCF:71Kb INV ESC using the Asb18 promoter as a viewpoint. The average Capture-C signals of the two replicates performed for each mESC line are shown around the Gbx2/Asb18 locus either individually (upper tracks) or after subtracting the WT Capture-C signals (lower tracks). The TAD boundary containing the three CTCF sites is highlighted in*

*light blue. The red arrows indicate the 71 Kb inversion breakpoints. (**B**) The average Capture-C signals shown in (A) were measured around the Gbx2 SE (highlighted in yellow in (A); chr1:89858398-89889044 (mm10)) as well as within five different 30 Kb control regions (Ctrls) located within the Asb18 TAD (red asterisks in (A) indicate the midpoint of the following controls regions (mm10): chr1:90049577-90079577, chr1:90099577-90129577, chr1:90149577-90179577, chr1:90199577-90229577, chr1:9024957-90279577). Capture-C signals are shown for the Δ3XCTCF, 71Kb INV and Δ3XCTCF:71Kb INV ESC as log2 fold-changes with respect to WT ESC.*

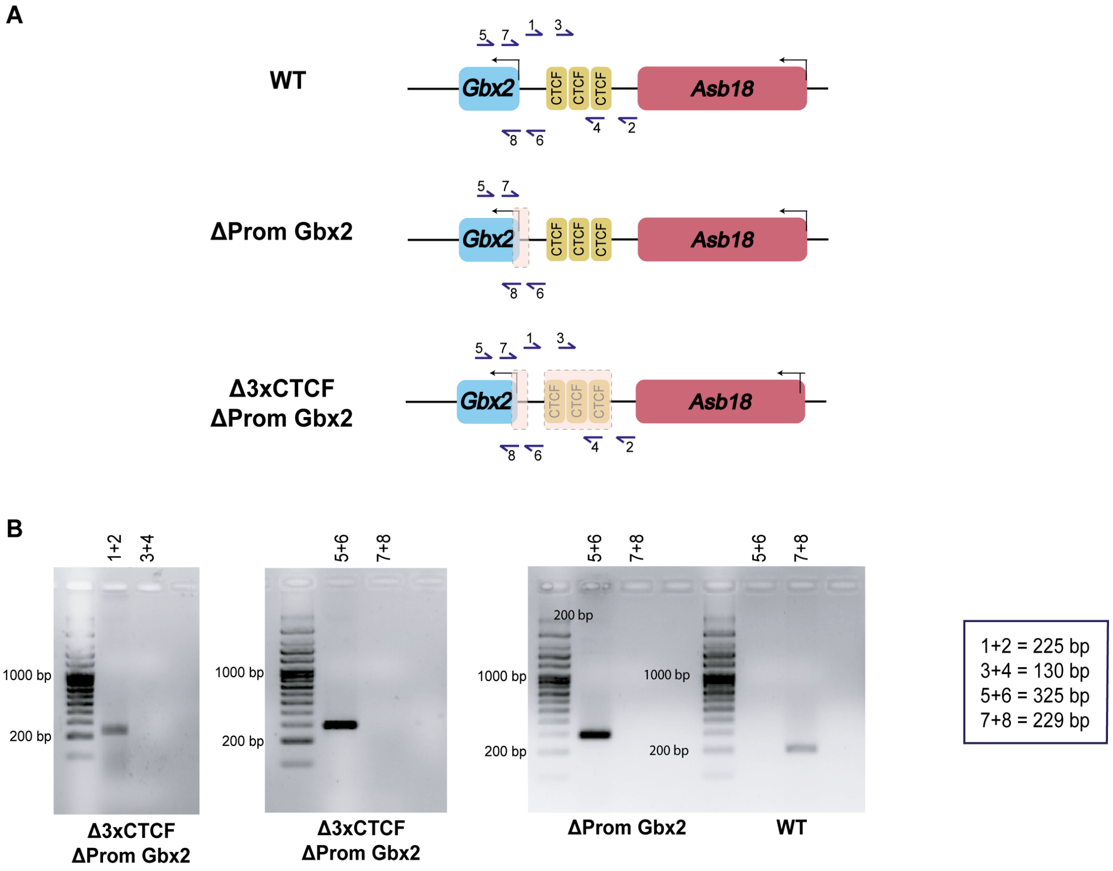


***Supplementary Fig. 9: Genotyping of the ΔPromGbx2 and Δ3XCTCF:ΔPromGbx2 re-arrangements generated at the Gbx2/Asb18 locus. (A)*** *Graphical overview of the PCR-based strategy used to genotype the ΔPromGbx2 and Δ3XCTCF:ΔPromGbx2 deletions described in Fig 4. The horizontal arrows and accompanying numbers represent PCR primers.* ***(B)*** *Representative PCR genotyping results obtained for ESC lines that were either WT or homozygous for the ΔPromGbx2 and Δ3XCTCF:ΔPromGbx2 deletions using the indicated primer pair combinations. The expected sizes of the amplicons obtained with each PCR primer combination are shown at the bottom right corner.*

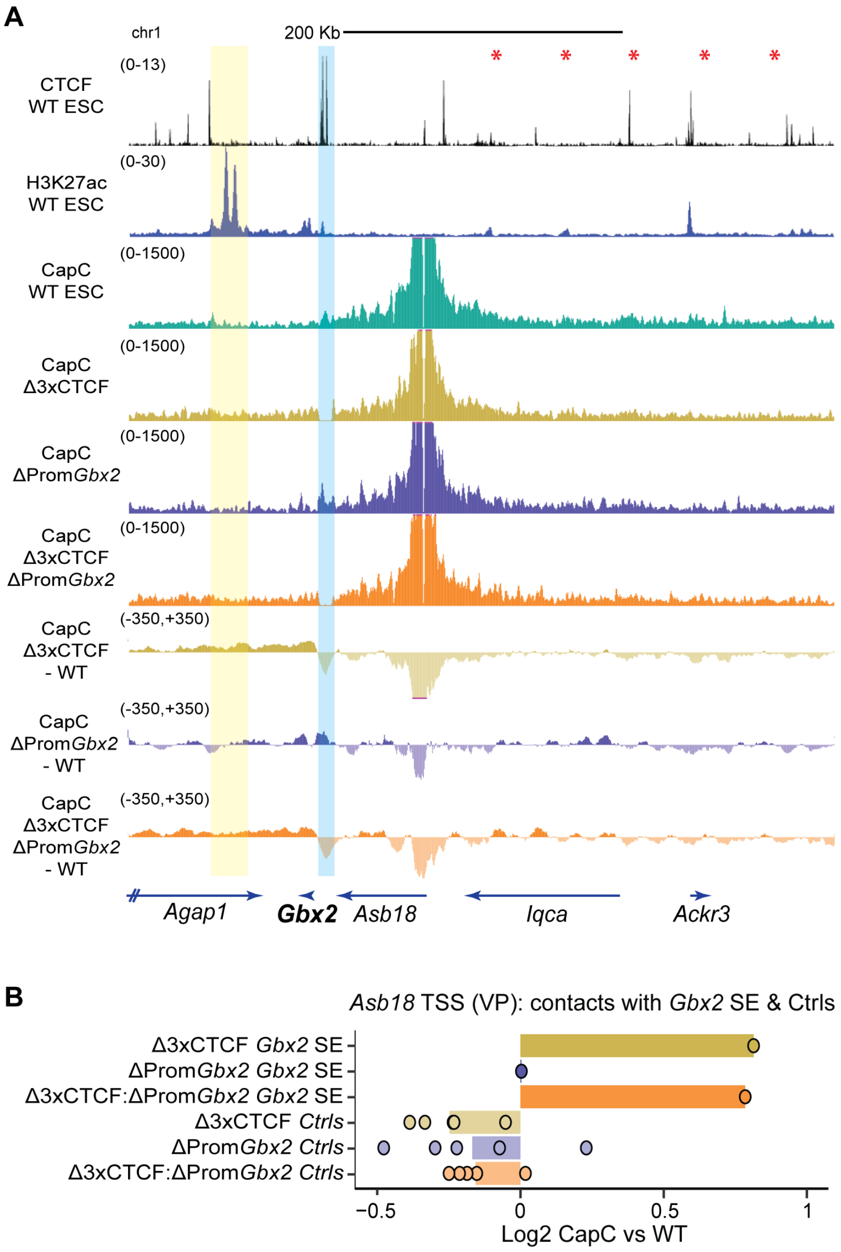


***Supplementary Fig. 10: Capture-C experiments in Δ3XCTCF, ΔPromGbx2 and Δ3XCTCF:ΔPromGbx2 mESC using the Asb18 promoter as a viewpoint. (A)*** *Capture-C experiments were performed as two biological replicates in WT, Δ3XCTCF, ΔPromGbx2 and Δ3XCTCF:ΔPromGbx2 ESC using the Asb18 promoter as a viewpoint. The average Capture-C signals of two replicates for each mESC line are shown around the Gbx2/Asb18 locus either individually (upper tracks) or after subtracting the WT Capture-C signals (lower tracks). The TAD boundary containing the three CTCF sites is highlighted in light blue. (**B**) The average Capture-C signals shown in (A) were measured around the Gbx2 SE (highlighted in yellow in (A); chr1:89858398-89889044 (mm10)) as well as within five different 30 Kb control regions (Ctrls) located within the Asb18 TAD (red asterisks in (A) indicate the midpoint of the same control regions as in Fig S8B). Capture-C signals are shown for the Δ3XCTCF, ΔPromGbx2 and Δ3XCTCF:ΔPromGbx2 ESC as log2 fold-changes with respect to WT ESC.*

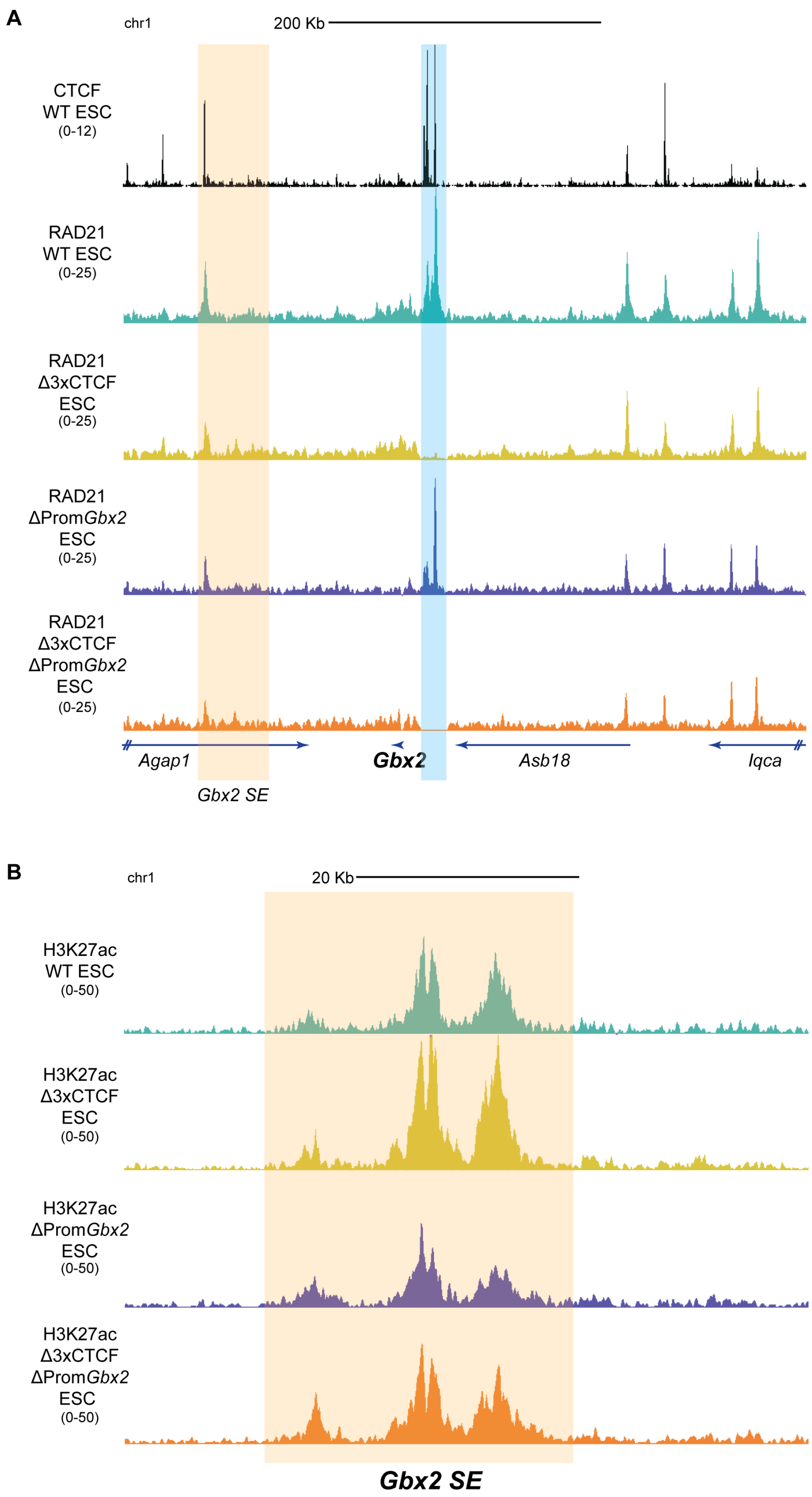


***Supplementary Fig. 11: RAD21 and H3K27ac profiles in ESC with genomic re-arrangements within the Gbx2/Asb18 locus. (A-B)*** *ChIP-seq profiles for (A) RAD21 and (B) H3K27ac are shown in ESC that are either WT or homozygous for the Δ3XCTCF, ΔPromGbx2 or Δ3XCTCF:ΔPromGbx2 deletions. In (A), CTCF ChiP-seq profiles generated in WT mESC* [102] *are also shown. The Gbx2 SE is highlighted in orange and the three CTCF sites deleted within the Gbx2/Asb18 TAD boundary are highlighted in blue.*

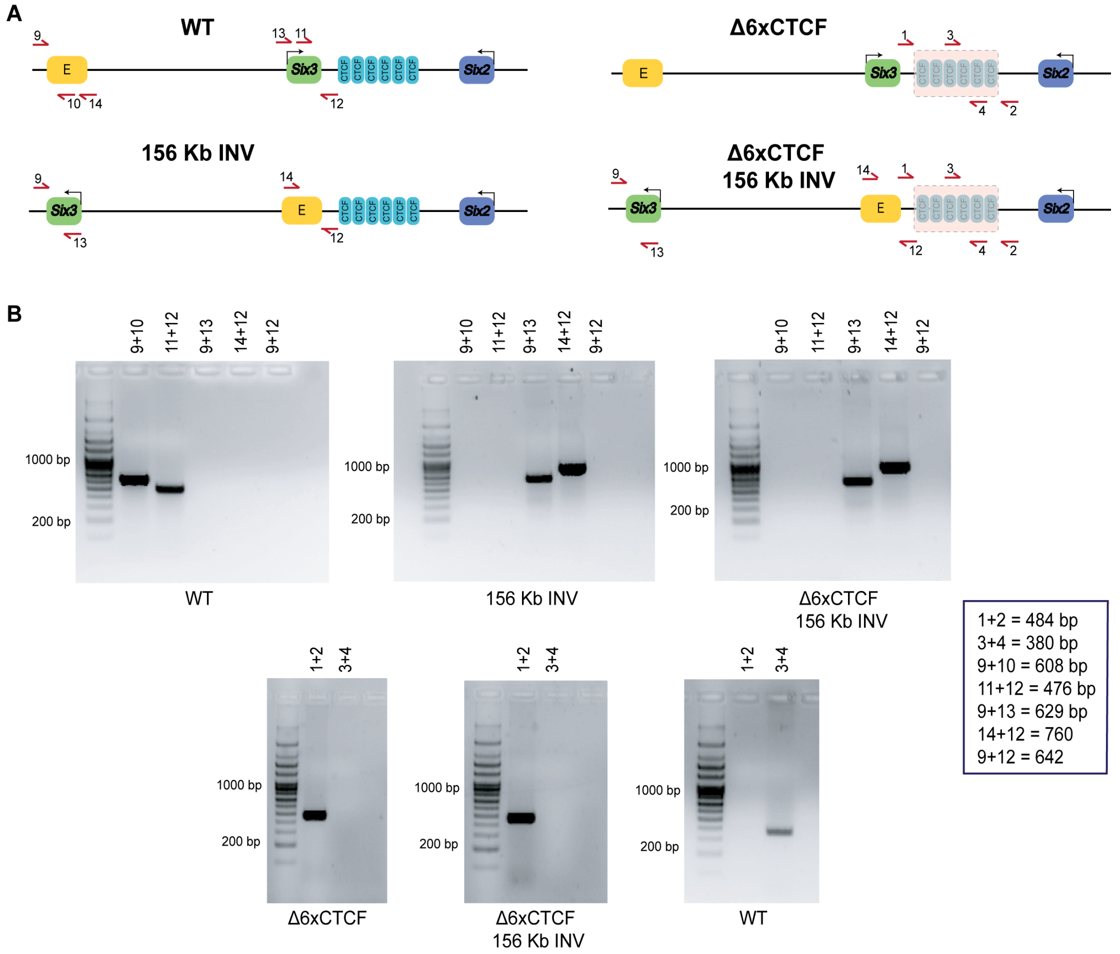


***Supplementary Fig. 12: Genotyping of the Δ6XCTCF, 156Kb INV and Δ6XCTCF:156Kb INV re-arrangements generated at the Six3/Six2 locus. (A)*** *Graphical overview of the PCR-based strategy used to genotype the Δ6XCTCF, 156Kb INV and Δ3XCTCF:156Kb INV genomic re-arrangements described in Fig 5. The horizontal arrows and accompanying numbers represent PCR primers.* ***(B)*** *Representative PCR genotyping results obtained for ESC lines that were either WT or homozygous for the 6XCTCF, 156Kb INV and Δ6XCTCF:156Kb INV genomic re-arrangements using the indicated primer pair combinations. The expected sizes of the amplicons obtained with each PCR primer combination are shown at the bottom right corner.*

.

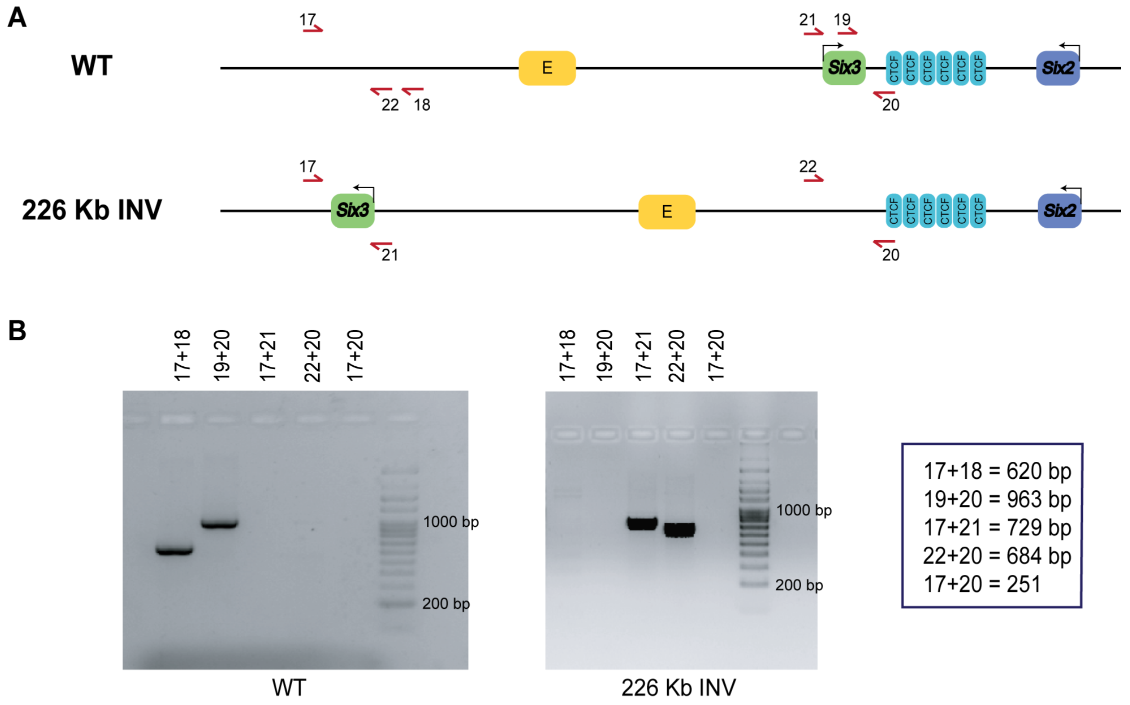


***Supplementary Fig. 13: Genotyping of the 226Kb INV ESC lines. (A)*** *Graphical overview of the PCR-based strategy used to genotype the 226 Kb INV inversion described in Fig 5. The horizontal arrows and accompanying numbers represent PCR primers.* ***(B)*** *Representative PCR genotyping results obtained for ESC lines that were either WT or homozygous for the 226 Kb INV inversion using the indicated primer pair combinations. The expected sizes of the amplicons obtained with each PCR primer combination are shown at the bottom right corner.*

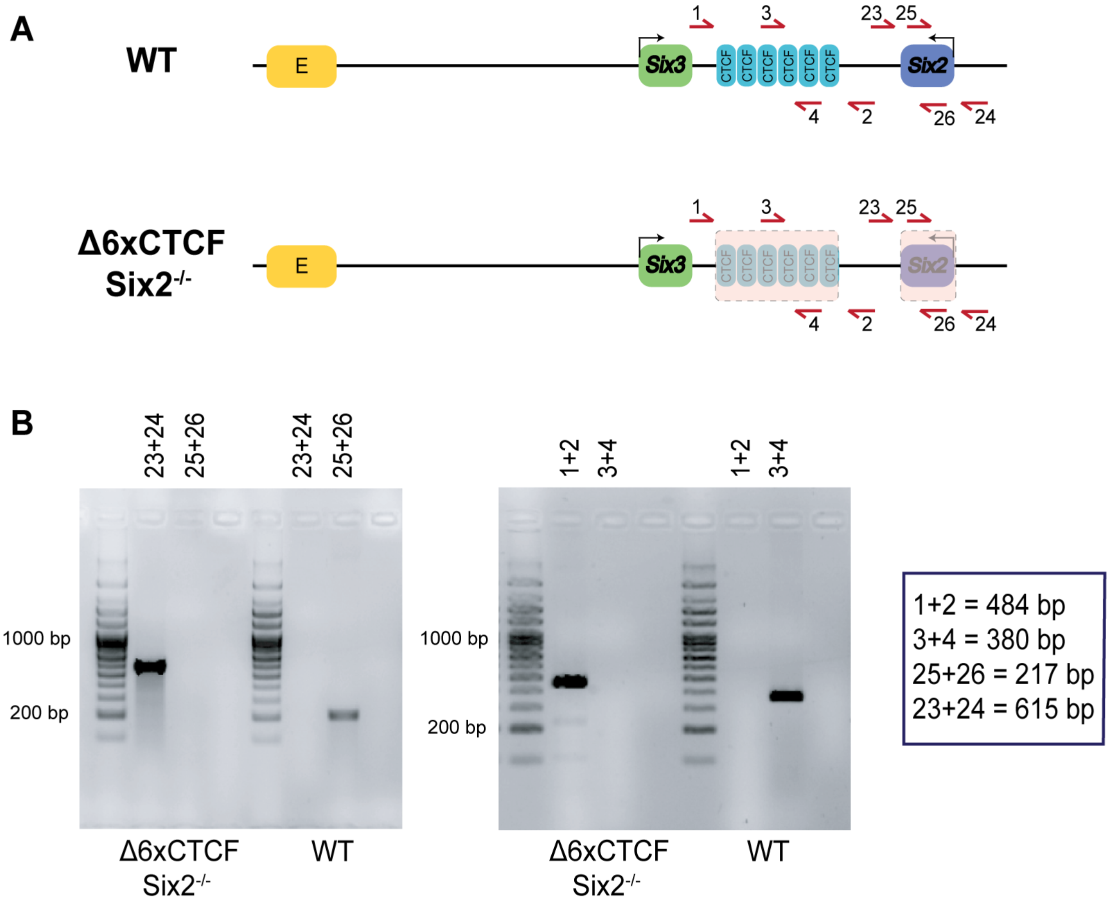


***Supplementary Fig. 14: Genotyping of the Δ6XCTCF:Six2$^{-/-}$ ESC lines. (A)*** *Graphical overview of the PCR-based strategy used to genotype the Δ6XCTCF:Six2$^{-/-}$ deletions described in Fig 5. The horizontal arrows and accompanying numbers represent PCR primers.* ***(B)*** *Representative PCR genotyping results obtained for ESC lines that were either WT or homozygous for the Δ6XCTCF:Six2$^{-/-}$ deletions using the indicated primer pair combinations. The expected sizes of the amplicons obtained with each PCR primer combination are shown at the bottom right corner.*

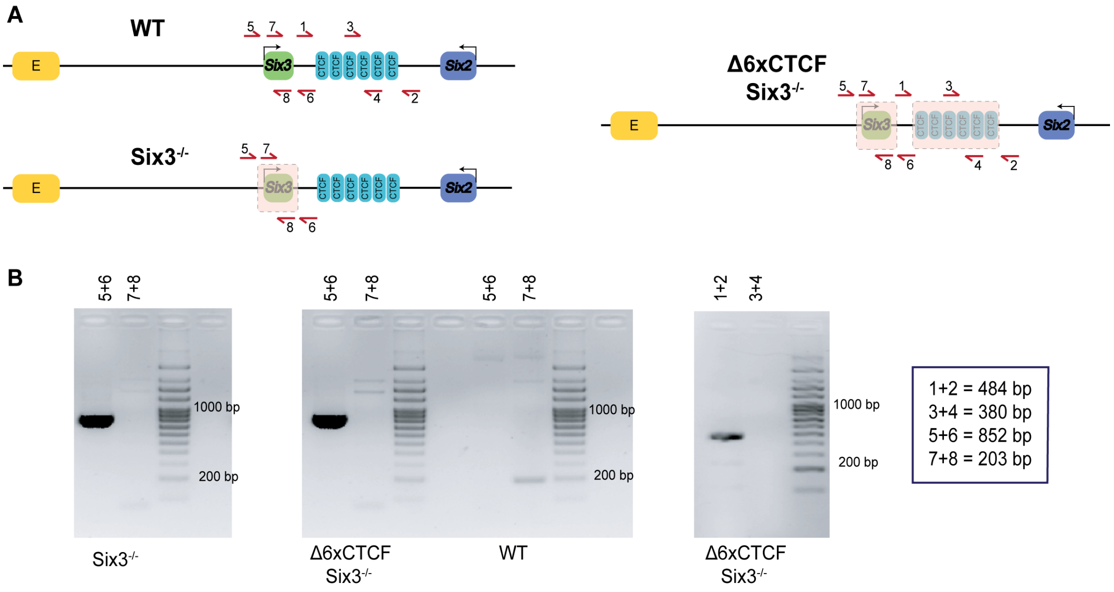


***Supplementary Fig. 15: Genotyping of the Six3$^{-/-}$ and Δ6XCTCF:Six3$^{-/-}$ deletions generated at the Six3/Six2 locus. (A)*** *Graphical overview of the PCR-based strategy used to genotype the Six3$^{-/-}$ and Δ6XCTCF:Six3$^{-/-}$ deletions described in Fig 6. The horizontal arrows and accompanying numbers represent PCR primers.* ***(B)*** *Representative PCR genotyping results obtained for ESC lines that were either WT or homozygous for the Six3$^{-/-}$ and Δ6XCTCF:Six3$^{-/-}$ deletions using the indicated primer pair combinations. The expected sizes of the amplicons obtained with each PCR primer combination are shown at the bottom right corner.*

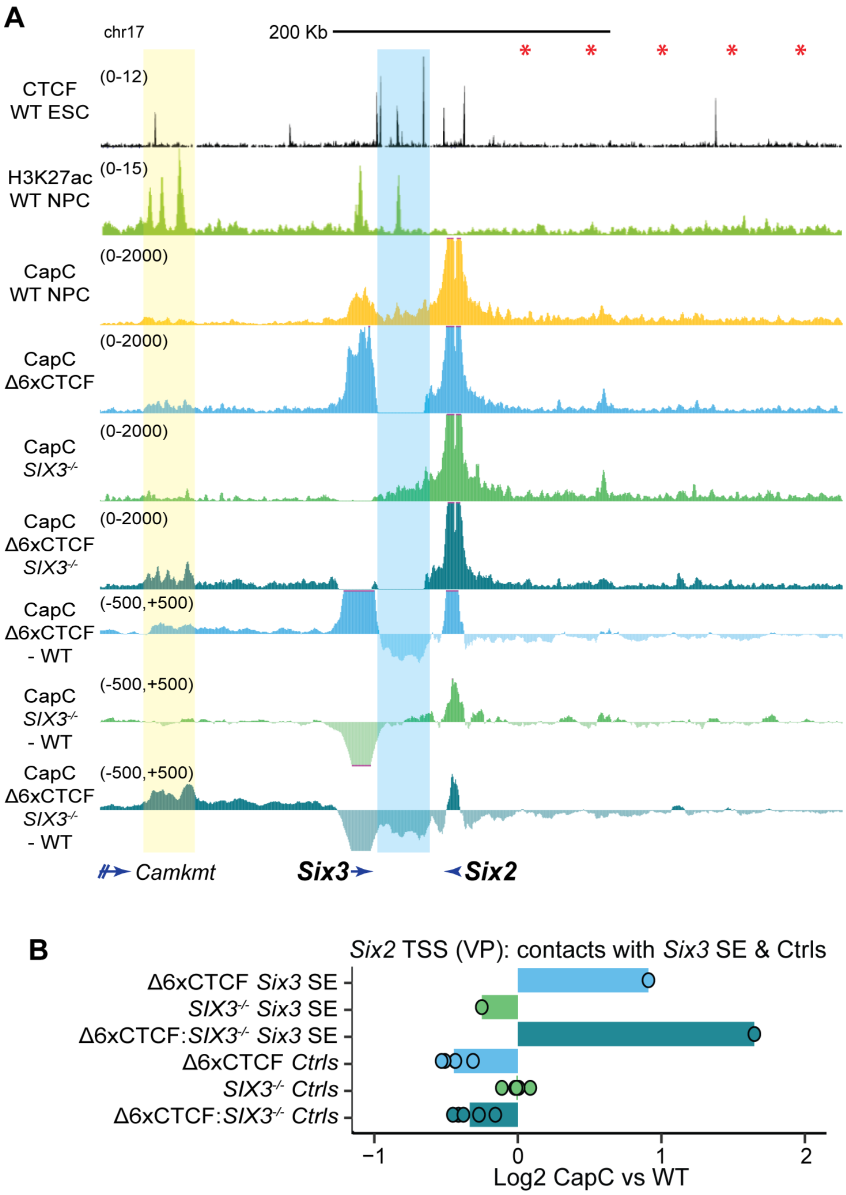


***Supplementary Fig. 16: Capture-C experiments in Δ6XCTCF, Six3$^{-/-}$ and Δ6XCTCF:Six3$^{-/-}$ NPC using the Six2 promoter as a viewpoint. (A)*** *Capture-C experiments were performed as two biological replicates in WT, Δ6XCTCF, Six3$^{-/-}$ and Δ6XCTCF:Six3$^{-/-}$ NPC using the Six2 promoter as a viewpoint. The average Capture-C signals of the two replicates performed for each cell line are shown around the Six3/Six2 locus either individually (upper tracks) or after subtracting the WT Capture-C signals (lower tracks). The six CTCF sites deleted within the Six3/Six2 TAD boundary are highlighted in blue. (**B**) The average Capture-C signals shown in (A) were measured around the Six3 SE (highlighted in yellow in (A); chr17:85453601-85491823 (mm10)) as well as within five different 30 Kb control regions (Ctrls) located within the Six2 TAD (red asterisks in (A) indicate the midpoint of the following controls regions (mm10): chr17:85723254-85753254, chr17:85773254-85803254, chr17:85823254-85853254, chr17:85873254-85903254, chr17:85923254-85953254). Capture-C signals are shown for the Δ6XCTCF, Six3$^{-/-}$ and Δ6XCTCF:Six3$^{-/-}$ NPC as log2 fold-changes with respect to WT NPC.*

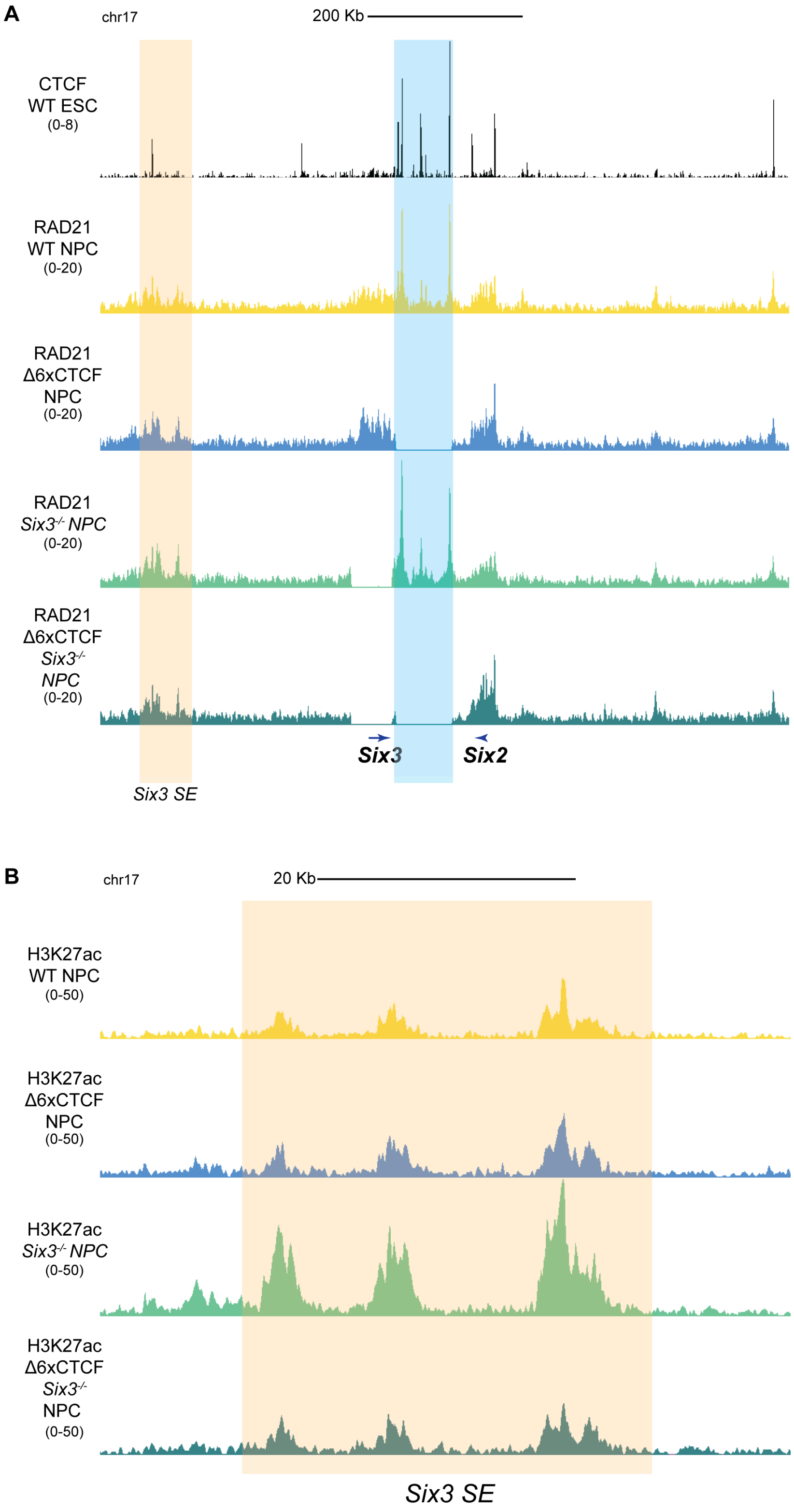


***Supplementary Fig. 17: RAD21 and H3K27ac profiles in NPC with genomic re-arrangements within the Six3/Six2 locus. (A-B)*** *ChIP-seq profiles for (A) RAD21 and (B) H3K27ac are shown in NPC that are either WT or homozygous for the Δ6XCTCF, Six3$^{-/-}$ or Δ6XCTCF:Six3$^{-/-}$ deletions. In (A), CTCF ChiP-seq profiles generated in WT mESC [102] are also shown. The Six3 SE is highlighted in orange and the six CTCF sites deleted within the Six3/Six2 TAD boundary are highlighted in blue.*

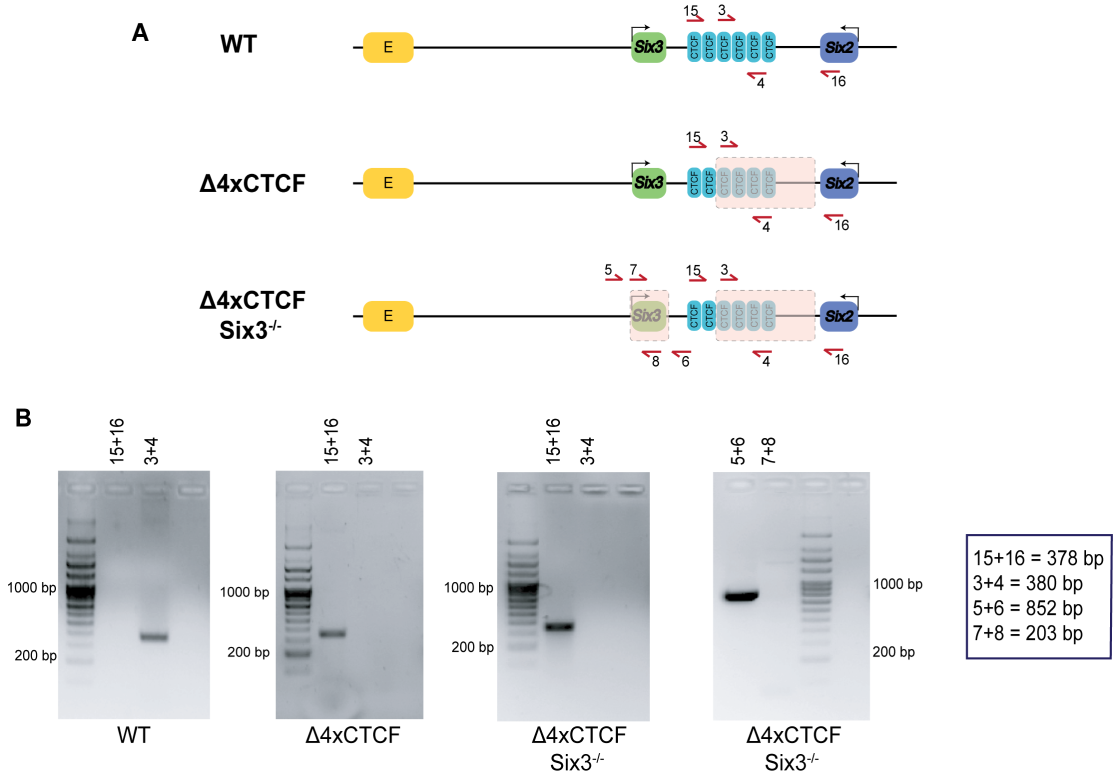


***Supplementary Fig. 18: Genotyping of the Δ4XCTCF and Δ4XCTCF:Six3$^{-/-}$ deletions generated at the Six3/Six2 locus. (A)*** *Graphical overview of the PCR-based strategy used to genotype the Δ4XCTCF and Δ4XCTCF:Six3$^{-/-}$ deletions described in Fig 6. The horizontal arrows and accompanying numbers represent PCR primers.* ***(B)*** *Representative PCR genotyping results obtained for ESC lines that were either WT or homozygous for the Δ4XCTCF and Δ4XCTCF:Six3$^{-/-}$ deletions using the indicated primer pair combinations. The expected sizes of the amplicons obtained with each PCR primer combination are shown at the bottom right corner.*
